# Nanophotonic DyeCycling delivers single-molecule FRET beyond photobleaching to identify heterogenous dynamics in DNA and protein systems

**DOI:** 10.64898/2026.09.23.753919

**Authors:** Benjamin Vermeer, Dong Hoon Shin, Alexander Vogel, Fabian Zundel, Sabina Caneva, Sonja Schmid

## Abstract

With its unique spatiotemporal resolution at the single-molecule level, Förster resonance energy transfer (FRET) is a powerful tool for investigating biomolecular conformational dynamics and function. However, early photobleaching remains a key limitation restricting the achievable observation time and therefore the information gain and application range of single-molecule FRET. Here, we establish nanophotonic DyeCycling, which overcomes photobleaching by reversible fluorophore binding and efficient background suppression using zero-mode waveguides. Nanoscale conformational changes are sensitively resolved from milliseconds up to the hour range, enabling robust kinetic analyses at the single-molecule level. As a result, outlier molecules reflecting static heterogeneity and time-dependent kinetics within individual molecules reflecting dynamic heterogeneity are reliably resolved. The versatility of DyeCycling is demonstrated using DNA and protein systems. Altogether, nanophotonic DyeCycling extends smFRET observations beyond the conventional photobleaching limit, revealing previously inaccessible kinetic effects and providing deeper insight into biomolecular systems with multiple states and rates.

---

Since their emergence, single-molecule biophysics experiments have resolved biological questions inaccessible to ensemble methods^1–4^. Their key advantage is the ability to identify molecular heterogeneity and to monitor dynamic changes in individual molecules, which led to a string of breakthroughs in biology. Well-known early examples include electrophysiology experiments revealing ion channel gating^5^, and the direct observation of the rotating ATP-synthase by fluorescence microscopy^6^, followed by many other experiments^7–11^. Among today’s single-molecule techniques, single-molecule FRET (hereafter smFRET) stands out due to its combined spatiotemporal resolution enabling researchers to follow the progression of a single biomolecule through distinct conformations along a defined reaction coordinate (**Fig. 1a**). This coordinate is set by two fluorescent probes that are chemically coupled to specific points of interest within the molecule under study. One probe serves as energy donor and the other as red-shifted acceptor for the strongly distance-dependent Förster resonance energy transfer. At short nanoscale distances, efficient FRET results in predominantly red emission. At larger distances (≥8 nm), FRET is minimal or absent, and green emission dominates. Both red and green emissions of single fluorophores are sensitively detected as a function of time with dedicated microscopes, as described in detail in recent reviews^12,13^. This ability of smFRET, to convert nanoscale distance changes into optical signals that are detectable at the single-molecule level, has led to important findings across diverse biomolecular systems^12–14^. Because fluorophores undergo irreversible photobleaching under laser excitation as a result of photodamage to their conjugated systems, considerable efforts have been made to reduce this limitation. In particular, the development of bright and stable fluorophores, including self-healing fluorophores^15–17^, triplet state quenchers^18–20^, and oxygen scavengers^21^, had a central role in resolving nanoscale effects in single surface-tethered molecules.

**Figure 1.**
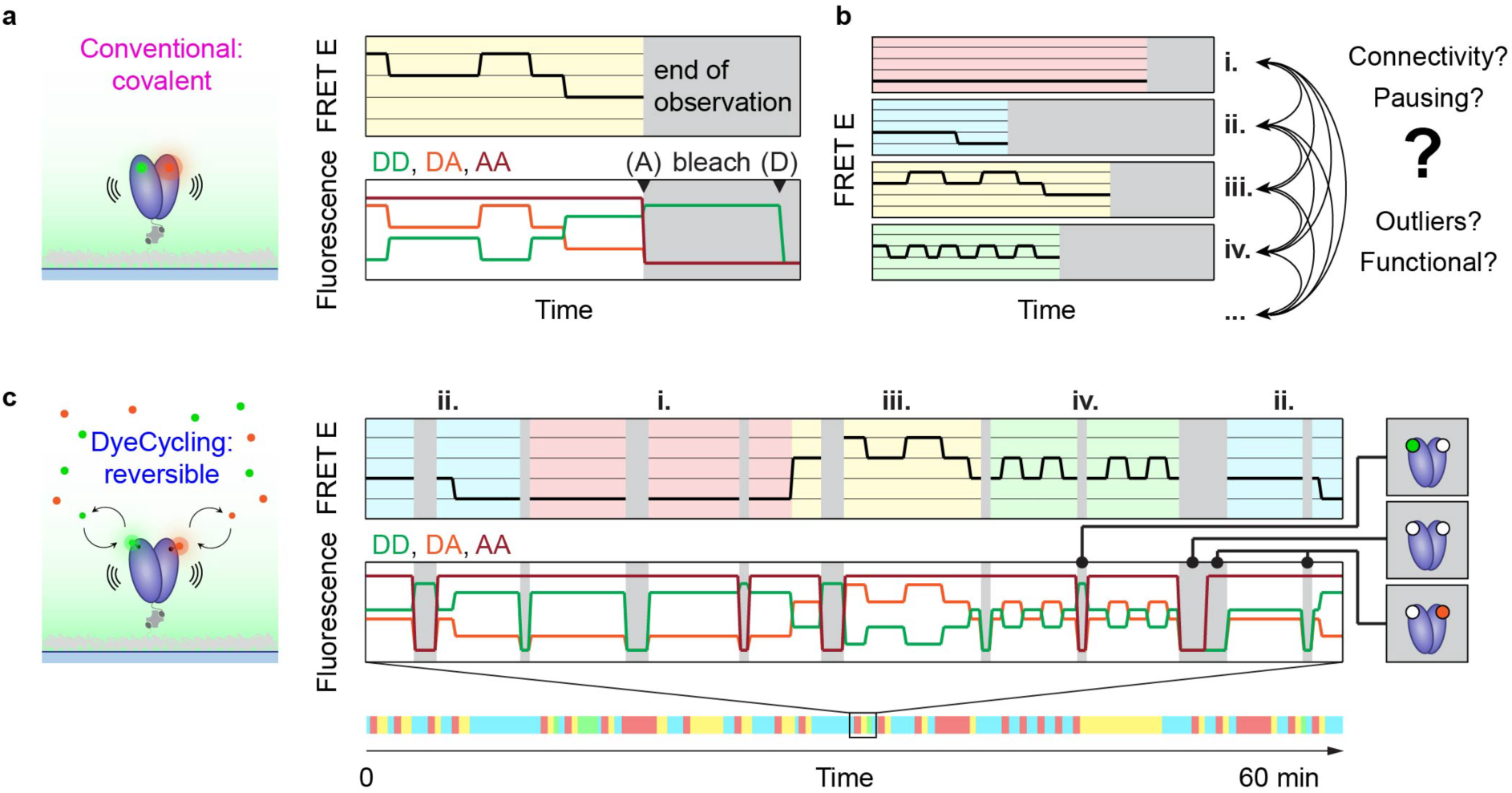
Conventional and DyeCycling-based smFRET experiments. (**a**) Conventional smFRET experiments to measure conformational changes in biomolecules are limited by photobleaching, leading to short observations. As the FRET fluorophores are irreversibly coupled to the biomolecule, the observation ends as soon as one of both fluorophores bleaches (grey shade). Acronyms: FRET E, FRET efficiency; D, donor; A, acceptor; DD, donor emission after donor excitation; DA, FRET-sensitized acceptor emission after donor excitation, AA, acceptor emission after acceptor excitation. (**b**) Due to photobleaching causing short observations, different data traces may show different kinetic behaviours (static, i; distinct dynamic behaviour ii, iii, iv), since the observation windows are too short to capture the full complexity of a biomolecule’s behaviour. This leads to several confounding effects, including uncertainty about the connectivity (i.e., the order in which the regimes occur in a functional cycle), whether static traces or rare events represent real biomolecular function or dysfunctional outliers. (**c**) DyeCycling overcomes the photobleaching limitation using reversibly bound fluorophores, providing long-term single-molecule observations to elucidate complex kinetic behaviours of biomolecules. Specifically, the connectivity (order) of the regimes (i) to (iv) is resolved and potential impurity outliers are identified. In this schematic example, static (i), distinct dynamic (ii, iii), and rare observations (iv) are detected to be real functional regimes that occur sequentially within a single molecule. Short intermittent gaps (grey) occur when one or both fluorophores are dissociated, as indicated on the right. Acronyms as in (a).

Still, photobleaching remains the main shortcoming of smFRET, in particular for kinetic trace analyses of complex biomolecular systems, which often involve multiple states connected by fast and slow transition rates. The limitation arises because three factors are coupled in a trade-off: the maximum achievable single-molecule observation time, time resolution, and signal-to-noise ratio all compete for a finite photon budget constrained by photobleaching. As a result, smFRET time series from single surface-tethered molecules span only 2-3 orders of magnitude in temporal bandwidth, measured from the shortest to the longest detectable interval (Fig. 1 of Ref ^22^). These bandwidths are substantially narrower than those of other single-molecule techniques (e.g. high-speed atomic force microscopy^23^, magnetic^24^ or optical tweezers^25^, nanopore detection^26^). There are several consequences of this limitation. First, the information contained in a single bleaching-limited smFRET trace is usually insufficient to make credible interpretations on molecular properties, in particular if more than two states and disparate kinetic rates are involved (illustrated in **Fig. 1a,b**). Therefore, many single-molecule traces are typically pooled together to perform an ensemble-averaged analysis (e.g., using FRET histograms and dwell-time distributions). However, the need for ensemble pooling negates a central benefit of single-molecule experiments, namely the unique ability to compare individual molecules and discover inter-molecular heterogeneities and co-existing sub-populations^24,27–29^. As a result, it is difficult to identify potentially damaged outlier molecules, mislabelled impurities, and also functional differences, for example, due to enzymatic or non-enzymatic post-translational modifications (e.g. spontaneous deamidation of asparagine and glutamine residues^30,31^). Next, even *within* individual biomolecules varying kinetic regimes can be observed on timescales beyond photobleaching. Literature examples include long-term pausing of fast RNA polymerases^24^, slow switching between fast exonuclease digestion rates^28^, sampling of heterogenous conformational ensembles by multi-domain chaperones^32–34^, and rare and decisive proline isomerizations next to fast conformer transitions^29^. In conventional, bleaching-limited smFRET experiments, such common multistate behaviour results in smFRET traces showing contrasting patterns illustrated in **Fig. 1b** – including so-called static traces (case i) and distinct types of dynamic traces (cases ii, iii, iv)^35–40^ – while the complete kinetic series remains inaccessible due to photobleaching. Moreover, whether these fragmented observations arise from multiple distinct molecular species (static heterogeneity) or from varying kinetic regimes of one single species (dynamic heterogeneity) generally remains unresolved. The dimensionality reduction from a 3D reality projected on the 1D reaction coordinate of smFRET further complicates this question. Taken together, it is clear that the well-known hierarchy of biomolecular dynamics^41^ spans broad timescales from <10^-6^ s to > 10^3^ s with short-lived events (≤ milliseconds; i.e. low energy barriers) to rate-limiting long-lasting events (minutes, hours; i.e. high energy barriers), and fully covering them would require a correspondingly broad temporal bandwidth, which bleaching-limited smFRET does currently not provide. This motivates the development of methods that can overcome the photobleaching limitation of smFRET to achieve long-term observations without sacrificing time resolution or signal-to-noise ratio, and thus better match the broad-range dynamics of biomolecular systems.

Here, we establish nanophotonic DyeCycling for DNA and protein systems and resolve conformational dynamics with high sensitivity, from milliseconds to the hour range. In contrast to conventional smFRET using irreversibly attached fluorophores, DyeCycling employs reversibly bound fluorescent probes that are continuously replaced from solution, thus decoupling smFRET measurements from individual photobleaching events. In introducing the DyeCycling approach^22^, we were inspired by previous applications of reversible binding, most notably in super-resolution imaging^42,43^, single-particle tracking^44^, and in measuring multiple FRET pairs within one molecule^45^. Previous attempts to establish DyeCycling-like experiments were hampered by prohibitively high fluorescence backgrounds, limiting the resolution of nanoscale conformational changes^46^. Using novel nanophotonic structures, we overcome this problem and achieve near-complete background suppression even at high fluorophore concentrations needed for DyeCycling. The increase in signal-to-noise ratio and the total observation time achieved by nanophotonic DyeCycling enables us to reveal and quantify static and dynamic heterogeneity prevalent even in simple DNA-based systems. To demonstrate the broader applicability of the approach, we apply DyeCycling to the multi-domain protein Hsp90, establishing the compatibility of the technique with proteins. Altogether, we demonstrate that nanophotonic DyeCycling extends smFRET beyond the conventional photobleaching limit, enabling the long-term observation of conformational dynamics and the resolution of kinetic effects that were previously inaccessible.

## RESULTS

### The concept of DyeCycling

A conventional smFRET measurement ends with the bleaching of one of both fluorophores (**Fig. 1a,b**). Conversely, DyeCycling overcomes this barrier by continuously replacing the fluorophores with new ones from solution. **Figure 1c** explains the general scheme: the surface tethered biomolecule of interest features binding sites, where freely diffusing fluorescent probes bind and dissociate spontaneously and specifically at the attachment point on the biomolecule of interest. In the experiment, this leads to consecutive bound and unbound intervals of the FRET donor and acceptor, causing FRET-sensitive intervals, intermitted by short ‘blind gaps’, where one or both fluorophores are missing. Two points are therefore important for the realization of DyeCycling: (i) optimal probe binding kinetics causing minimal blind gaps, and (ii) efficient suppression of the background fluorescence of the probe solution, which are both detailed below. Once implemented, the presence and absence of each FRET fluorophore is determined using alternating laser excitation (ALEX)^47^ of the donor and acceptor, resulting in three recorded fluorescence intensities: directly excited donor (DD in **Fig. 1c**), FRET-sensitized acceptor (DA), and directly excited acceptor (FRET-independent, AA). Based on this 3-dimensional ALEX data, the blind gaps are easily identified as schematically illustrated on the right of **Fig. 1c**, and conformational dynamics between distinct FRET states can be resolved as usual from the FRET-sensitive intervals. We automated the identification of the conformational states and the blind gaps using Hidden Markov modelling (HMM) as frequently used in smFRET^48^. Specifically, we harnessed our pre-validated workflow, termed Single-Molecule Analysis of Complex Kinetic Sequences (SMACKS)^48,49^, which enables kinetic rate extraction also in the presence of blind gaps (see Methods). In brief, conventional smFRET provides bleaching-limited, short observations (**Fig. 1a**) that can be challenging to interpret, because individual molecules often exhibit distinct behaviours within these short observation times (**Fig. 1b**). By enabling smFRET beyond photobleaching, DyeCycling can detect biomolecular dynamics over long timescales (**Fig. 1c**) to resolve previously missed effects, such as static and dynamic heterogeneities.

### Design principles for reversible fluorescent probes

In principle, DyeCycling can be realized with various reversible chemistries, as previously proposed (reversible covalent bonds, host-guest or coordination chemistry, etc.)^22^. Here, we chose short single-stranded DNA oligomers (ssDNA) as specific docking sites for fluorescently labelled complementary ssDNA, whose dissociation rate is adjustable by ssDNA length and sequence. The goal for the DyeCycling experiment is to achieve (i) minimal unbound gaps and (ii) maximum bound time without suffering from photobleaching under the constraints set by the biomolecular dynamics under study. In short, this translates to two general design principles for the fluorescent probes: (i) a fastest possible binding rate and (ii) a slowest possible dissociation rate within the boundary set by the targeted dynamics. As an example, for the Holliday junction dynamics measured herein (∼1 – 4 s^-1^), rates of ∼1 s^-1^ for binding and ∼0.1 s^-1^ for dissociation would be ideal, given the bleaching rate of 0.06 s^-1^ found in conventional smFRET experiments with optimized laser power and time resolution (ALEX mode: 50 ms green, 50 ms red exposure). Indeed, a screen of several designs revealed that nine nucleotides with a 56% GC content yielded a dissociation rate of 0.13 s^-1^ (see Methods, **Table 1**). Next, since minimal unbound gaps are crucial for DyeCycling, the binding rate was increased by optimising ssDNA sequences to prevent self-complementarity^50^. In addition, utilizing high concentrations (µM) of freely diffusing fluorescent probes can minimize unbound times but is incompatible with single-molecule resolution in regular TIRF microscopy. Hence, for more efficient fluorescent background suppression – without complicating fluorescent probe design through fluorogenicity – we developed nanophotonic devices described in the next section. These devices enabled sensitive single-molecule resolution in micromolar solutions of fluorescent probes, resulting in a binding rate of ca. 1.3 s^-1^. Taken together, these binding and dissociation rates of the fluorescent probes offer a theoretical temporal coverage of 90% per single probe (i.e. 10% gaps) leading to a theoretical coverage of 81% for the complete FRET pair (i.e. 19% gaps).

**Table 1.**
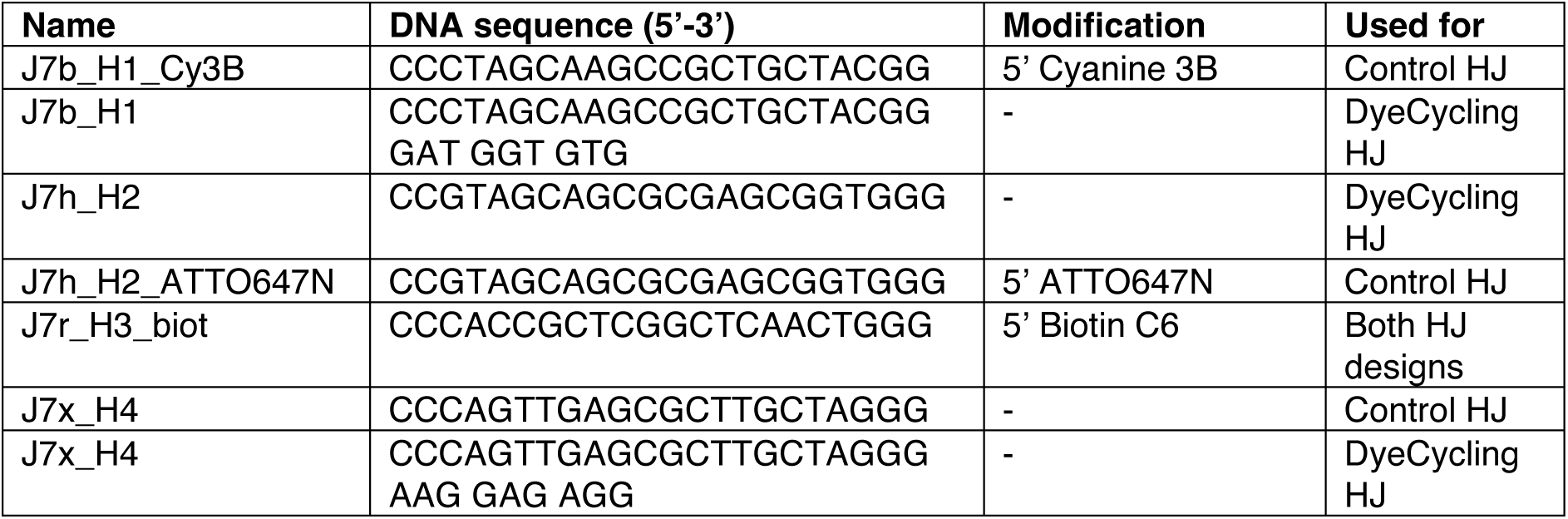
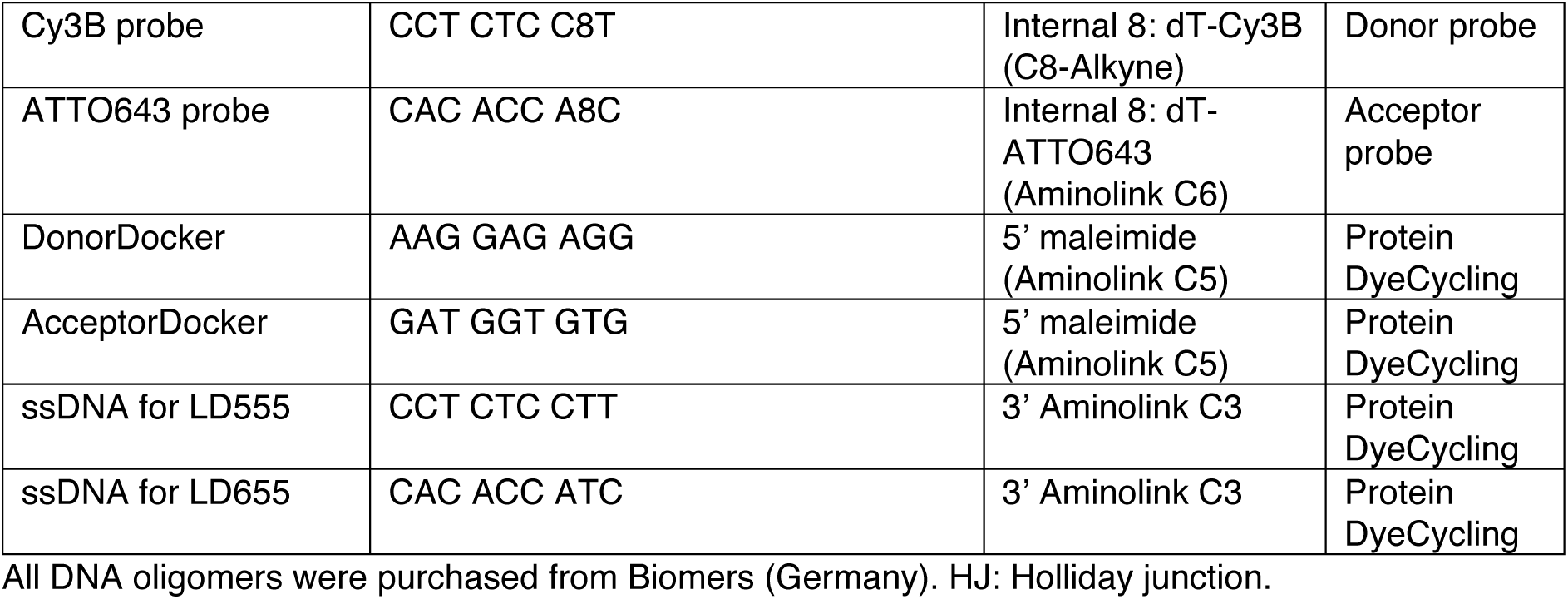
ssDNA oligomers used in this study.

### Nanophotonic suppression of background fluorescence

Efficient background suppression is essential for the realization of DyeCycling experiments, specifically to retain single-molecule resolution in micromolar fluorophore solutions enabling fast binding and short unbound gaps. To achieve this *without* restricting the choice of fluorophores and probe design, we opted for nanophotonics rather than fluorogenic strategies^46,51^. We chose zero-mode waveguides (ZMW), which are well-known nanophotonic devices consisting of nano-wells in a metal film^52–54^ (**Fig. 2a-c**). By blocking the propagation of light beyond the nano-well, they prevent the excitation of fluorophores outside of the nano-well, limiting the excitation volume down to zeptoliters^52^. Specifically for DyeCycling, we developed novel ZMWs optimized for the high-throughput detection of smFRET on surface-tethered molecules, maximising both signal-to-noise ratio and trace yield. The ZMW design parameters differ substantially from previously reported ZMWs, optimized for confocal smFRET detection^55,56^, nanopores^57^, imaging^54^, or sequencing^53,58^. In brief, we chose palladium (Pd) as the ZMW metal because of its superior properties compared to other common metals (Al, Au), including chemical resistance, undetectable autofluorescence, and favorable fabrication properties^57^ (see Methods). We then fabricated large ZMW arrays containing 10’000 apertures per field-of-view (**Fig. S1**) by focused ion beam (FIB) milling, achieving high precision and reproducibility of the design parameters described below. Before all measurements, the ZMWs were silanized and functionalized with polyethylene glycol for passivation and surface tethering via biotin-neutravidin coupling (Methods).

**Figure 2.**
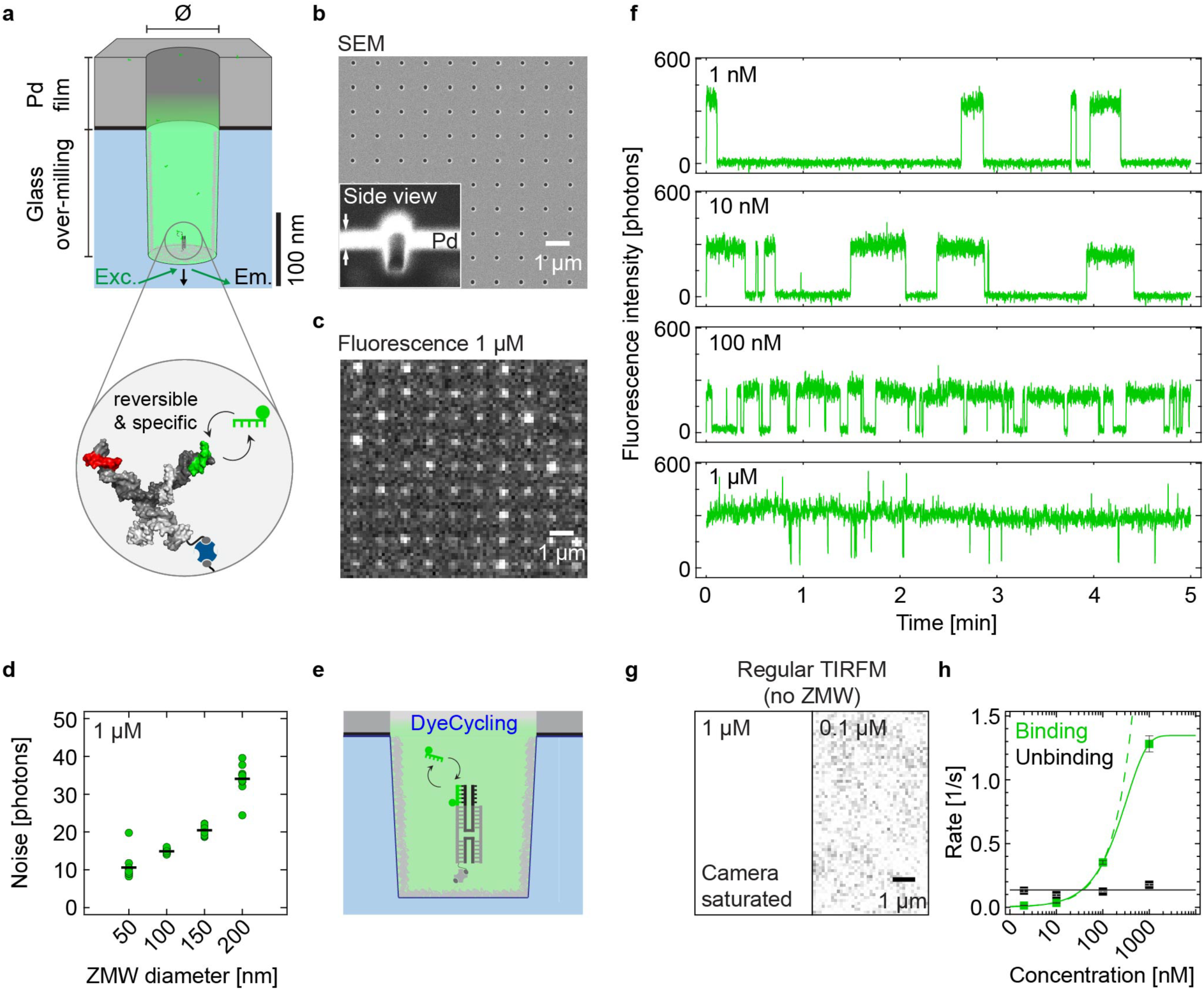
Efficient fluorescence background suppression with palladium zero-mode waveguides. (**a**) Schematic zero-mode waveguide (ZMW), a 100 nm diameter nano-well in a 100 nm palladium film on glass with 175 nm over-milling (see Methods). The zoom-in shows a surface-tethered Holliday junction with ssDNA overhangs for reversible binding of the fluorescent probe. (**b**) Scanning electron micrograph of a subset of a ZMW array. Inset: cross-section view of a single waveguide. White arrows indicate the 100 nm metal layer. (**c**) Fluorescence micrograph of the experiment in (a): a subset of a ZMW array with 1 µM fluorescent probes reversibly binding to surface-tethered biomolecules. (**d**) ZMW-diameter dependence of background noise (standard deviation) caused by 1 µM fluorescent probes. Data from ten ZMW (green, standard deviation of intensity) and their mean (black bar) are shown per diameter. See also **Fig. S3,4**. (**e**) Illustrated single-colour DyeCycling with a Holliday junction (not to scale), as used for (f,g). (**f**) Single-colour DyeCycling recordings at increasing probe concentrations as specified show minimal unbound gaps at 1 µM concentration (time resolution: 100 ms). (**g**) Comparison with a TIRF micrograph of a standard coverslip with 1 µM and 100 nM fluorescent probes in solution as indicated. Single-molecule resolution is not possible. (**h**) Concentration dependence of the kinetic rates of single-colour DyeCycling (binding/unbinding): means and 95% confidence intervals are plotted with logarithmic x-axis. The dissociation rates (black) remain constant, while the binding rates scale linearly (dashed green line) until bound-time saturation (green line). Fluorescent probe used: Cy3B probe (**Table 1**).

First, we screened various depths of over-milling into the glass (**Fig. 2a,b**) and found best high-quality trace yields with 150-180 nm over-milling combined with TIR illumination, which we used for all further experiments. In addition to favourable excitation field enhancement previously described for fluorescence imaging^54^, we attribute the benefit of over-milling for our smFRET measurements to the (on average) larger separation of the surface-tethered molecule from the metal, reducing fluorophore-metal interactions, such as fluorescence quenching or other interference with FRET^55,59–62^. Indeed, in contrast to previous results on confocal smFRET in different ZMW^55^, these newly designed ZMW do not measurably alter the FRET process, yielding FRET efficiencies consistent with coverslips-based surface-tethered experiments (**Fig. S2**). Next, we systematically optimized the ZMW diameter. We fabricated ZMWs with diameters varying from 50 to 200 nm and assessed their background suppression capacity, quantified by the residual background intensity and noise level in the presence of fluorescent probes at increasing concentrations. **Figure 2d** shows the expected increase in background noise as a function of ZMW diameter (see **Fig. S3,4** for more donor and acceptor data). We then assessed the single-molecule resolution of these ZMW in the presence of fluorescent background, using surface-tethered Holliday junctions with single-strand overhangs as docking strands for the fluorescent probes (illustrated in **Fig. 2e**). By testing increasing probe concentration, we identified a ZMW diameter of 100 nm as the optimum, balancing efficient background suppression (smaller being better, **Fig. 2d**) with good trace yield (larger being better). The resulting trace yield was 30-50% of that obtained with coverslip-based smFRET measurements (**Table S1**). **Figure 2f** shows such single-molecule fluorescence traces, revealing reversible probe-binding events measured at nanomolar to micromolar concentrations. Notably, the achieved high signal-to-noise ratio provides single-molecule resolution even at micromolar probe concentrations, where TIRF microscopy reaches detector saturation (**Fig. 2g**). As expected, the binding rate of the fluorescent probe increases linearly with concentration until saturation at 1 µM (**Fig. 2h**), while the dissociation rate remains constant.

In summary, we developed large, chemically inert, and reusable palladium ZMW arrays with excellent background suppression capacity, enabling low-noise single-molecule detection even in micromolar fluorophore solutions. In the context of DyeCycling, this allows reducing unbound gaps to durations approaching the bound-time saturation limit, as visible in the fluorescence traces and the binding and dissociation rates in **Figures 2f,h**, while the background noise stays very low (**Fig. 2h, Fig. S3,4**).

### Hour-long DyeCycling resolves nanoscale conformational changes by smFRET

Equipped with the two key ingredients of the DyeCycling technique – reversibly binding fluorescent probes and efficient background suppression – we proceeded to resolve the nanoscale conformational changes of single DNA Holliday junctions, as a well-studied dynamic test system^63,64^. Nanophotonic DyeCycling resolved these nanoscale rearrangements with comparable sensitivity to bleaching-limited smFRET (**Fig. 3a-c**), while providing substantially extended observation times. **Figure 3d** compares the FRET histograms obtained from single traces in each mode, demonstrating that DyeCycling provides consistent, well resolved FRET peaks while greatly increasing the amount of data detectable from a single molecule.

**Figure 3.**
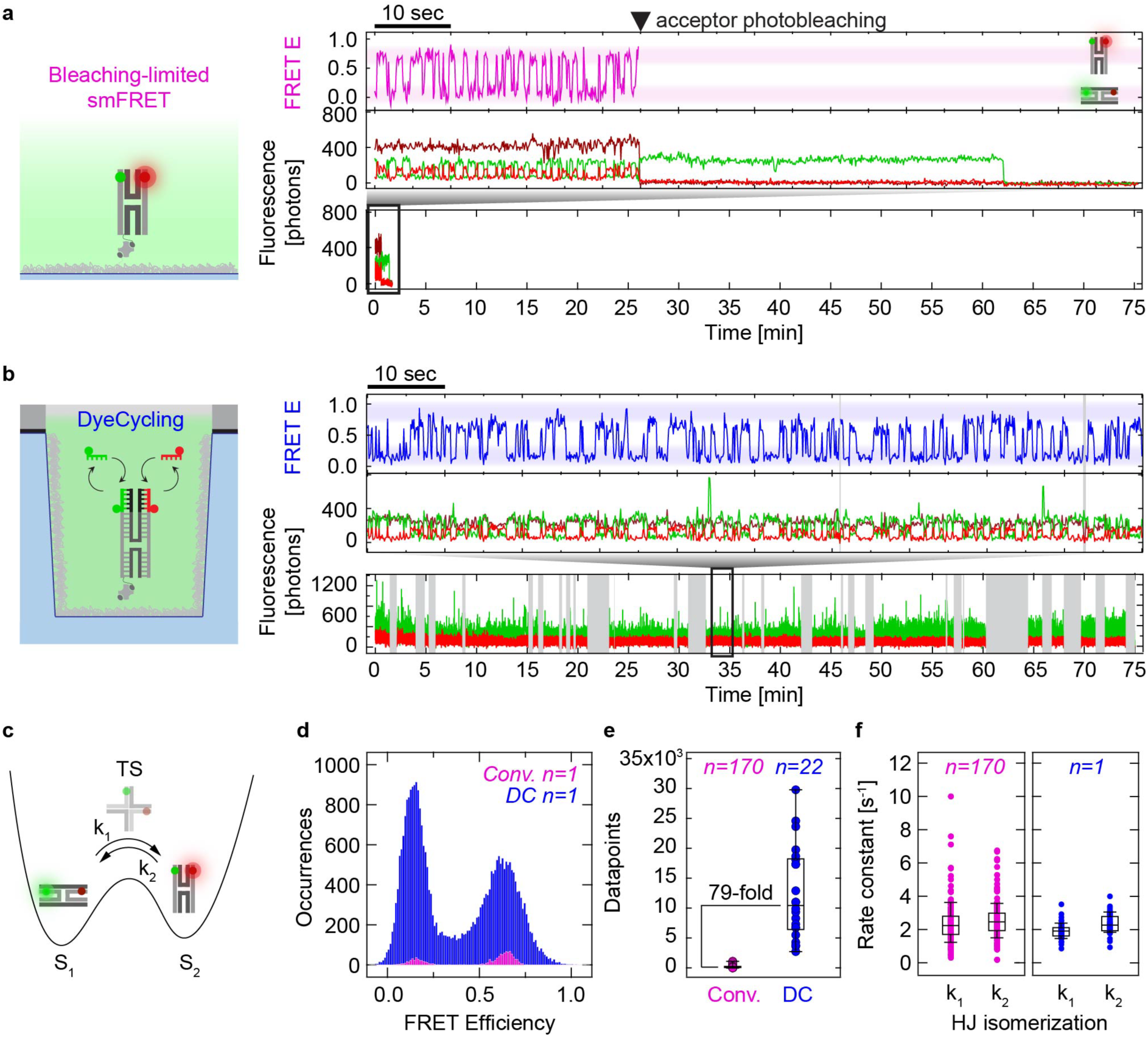
Comparison of conformational dynamics resolved by bleaching-limited smFRET vs. hour-long DyeCycling. (**a**) DNA Holliday junction dynamics resolved by bleaching-limited smFRET. The observation lasts for 35 seconds, as visible from the zoom-view (FRET E: FRET efficiency). Sampling rate: 10 Hz (50 ms green, 50 ms red excitation). Green: donor fluorescence; red: FRET-sensitized acceptor fluorescence; dark red: directly excited acceptor fluorescence. Right: cartoons of the low- and high-FRET conformations of the Holliday junction. (**b**) DNA Holliday junction dynamics resolved by nanophotonic DyeCycling in Pd ZMW (cartoon not to scale). The observation lasts for 75 min. Sampling rate: 10 Hz (50 ms green, 50 ms red excitation). Grey shading: (partially) unbound gap intervals. A FRET E zoom-view shows the well resolved conformational dynamics. The directly excited acceptor intensity was omitted for clarity; it is provided in **Fig. S9**. (**c**) Schematic energy landscape of the Holliday junction including equilibrium states S_1_, S_2_, transition state (TS) and kinetic rates k_1_ and k_2_. (**d**) Comparison of the FRET efficiency histograms of a single molecule measured by either conventional bleaching-limited smFRET (pink, covalently labelled) or DyeCycling (blue). Comparison of the longest trace of each kind. (**e**) Comparison of the number of datapoints measured per molecule by conventional bleaching-limited smFRET (pink, median: 132) versus DyeCycling (blue, median: 10442, excluding (partially) unbound gaps). Box plots show percentiles (0, 25, 50 (median), 75, and 100). (**f**) Comparison of the kinetic rates indicated in (c), obtained by bleaching-limited smFRET (pink, n=170 Holliday junction molecules) versus DyeCycling (blue, n=1 molecule split into 70-second segments). Box plots show percentiles (10, 25, 50 (median), 75, 90). (a-f) Consistent buffer and scavenger conditions, as well as sampling rates (10 Hz, 50 ms green, 50 ms red excitation) were used throughout **Fig. 3,4** and **Fig. S6a** (see Methods). Fluorescent probes used: Cy3B and ATTO643 probes (**Table 1**). FRET windows < 1s were discarded from the DyeCycling data (see Methods).

The conventional smFRET experiment ends by photobleaching (after 35s in **Fig. 3a**), causing short exponentially distributed observation times (see **Fig. S5** for a whole-dataset plot). Conversely, the DyeCycling experiment is not limited by a single bleach event and can run over an hour, albeit with intermittent gaps. To identify the gap intervals automatically, we developed an analysis pipeline based on HMM using the SMACKS software^49^. In short, it uses all three fluorescence intensities measured by ALEX and classifies the gaps with dissociated donor and/or acceptor probes as individual states (see Methods, **Fig. S7,8** for details). Based on this 3-fold input data, gap identification worked reliably, preventing misinterpretations (**Fig. S8**). We realized this Holliday junction experiment with two different FRET pairs (see **Fig. S6a,b** for whole-dataset plots). For realization 1 using Cy3B/Atto643, the temporal coverage was 62% over the entire dataset (i.e., recording time minus gaps), and for realization 2 using self-healing LD555/LD655^17^, it was 68% (at slightly higher probe concentrations). Both values are below the theoretical maximum coverage of 81% predicted by the binding kinetics discussed above. We mainly attribute this result to photochemical effects, affecting the reversible binding capacity of the ssDNA docking strands as previously reported^65^, while the comparison of realization 1 and 2 shows that higher probe concentrations can still improve temporal coverage in the future. Nonetheless, compared with conventional smFRET, DyeCycling data provides a large information gain, quantified here by the number of FRET-sensitive datapoints detected per molecule resulting in a 79-fold median gain (**Fig. 3d**). As a result, the DyeCycling dataset (realization 1) contains 9-fold more data than the conventional dataset (272’348 vs. 30’590 datapoints), despite 8-fold fewer traces. These numbers convert to an average of 12’379 datapoints/molecule or 20.6 minutes total observation time for DyeCycling at 10 Hz sampling, compared to an average of 179 datapoints/molecule or 17.9 seconds observation time for the conventional smFRET experiment.

Next, we assessed the conformational kinetics resolved and quantified from DyeCycling data. Using the SMACKS-based analysis described above, the gaps of the DyeCycling data were filtered out and the rate constants were directly extracted from the transition matrix of the Hidden Markov model. Briefly, this pre-validated rate extraction approach^48,49^ avoids dwell-time segmentation and related data loss, since every datapoint counts (rather than every dwell time), regardless of whether it belongs to a transition or a static interval. **Fig. 3f** compares kinetic rates extracted from 170 bleaching-limited smFRET traces with those of a single DyeCycling trace (cut into 70-second segments), confirming consistent kinetics are extracted from conventional and DyeCycling data.

In summary, DyeCycling preserves the core strengths of conventional smFRET – clearly resolved conformational populations and consistent kinetics – while delivering more than an order of magnitude more information per molecule. This increase in information content finally enables single-molecule analysis of single-molecule FRET data, including FRET efficiencies and kinetics, uniquely complementing the ensemble averaging generally required in conventional smFRET. Although demonstrated here on the well-characterized Holliday junction as a validation system, these capabilities are especially relevant for more complex molecules such as ribozymes and proteins, for which DyeCycling was originally devised.

### Revealing static and dynamic heterogeneity

In the following, we demonstrate the benefits of the validated single-trace kinetic analysis enabled by the long DyeCycling traces. Conventionally, standard kinetic analyses involve ensemble pooling of bleaching-limited smFRET traces, from which one set of kinetic rates is obtained per dataset; here k_1_ = 2.2 ± 1.0 s^-1^ and k_2_ = 2.5 ± 1.0 s^-1^ (mean ± standard deviation, **Fig. 4a**). This is because short bleaching-limited traces generally provide insufficient statistics and fragmented observations of the underlying kinetics, observed as “static molecules” and varied “dynamic molecules” introduced in **Fig. 1b**. It therefore remains unresolved which of the varied observations originate from distinct molecular species (static heterogeneity, e.g. due to misfolding, mislabelled impurities, lasting interaction with the environment (sticking), post-translational modifications, etc.) and which represent distinct kinetic regimes intrinsic to the complex dynamic behaviour of a single species (dynamic heterogeneity^24,28,29,32–34^, i.e. due to complex energy landscapes with different minima separated by high energy barriers). Because these cases cannot be distinguished by conventional smFRET, the most common workflow is to pool all traces together and extract ensemble-averaged kinetic rates.

**Figure 4.**
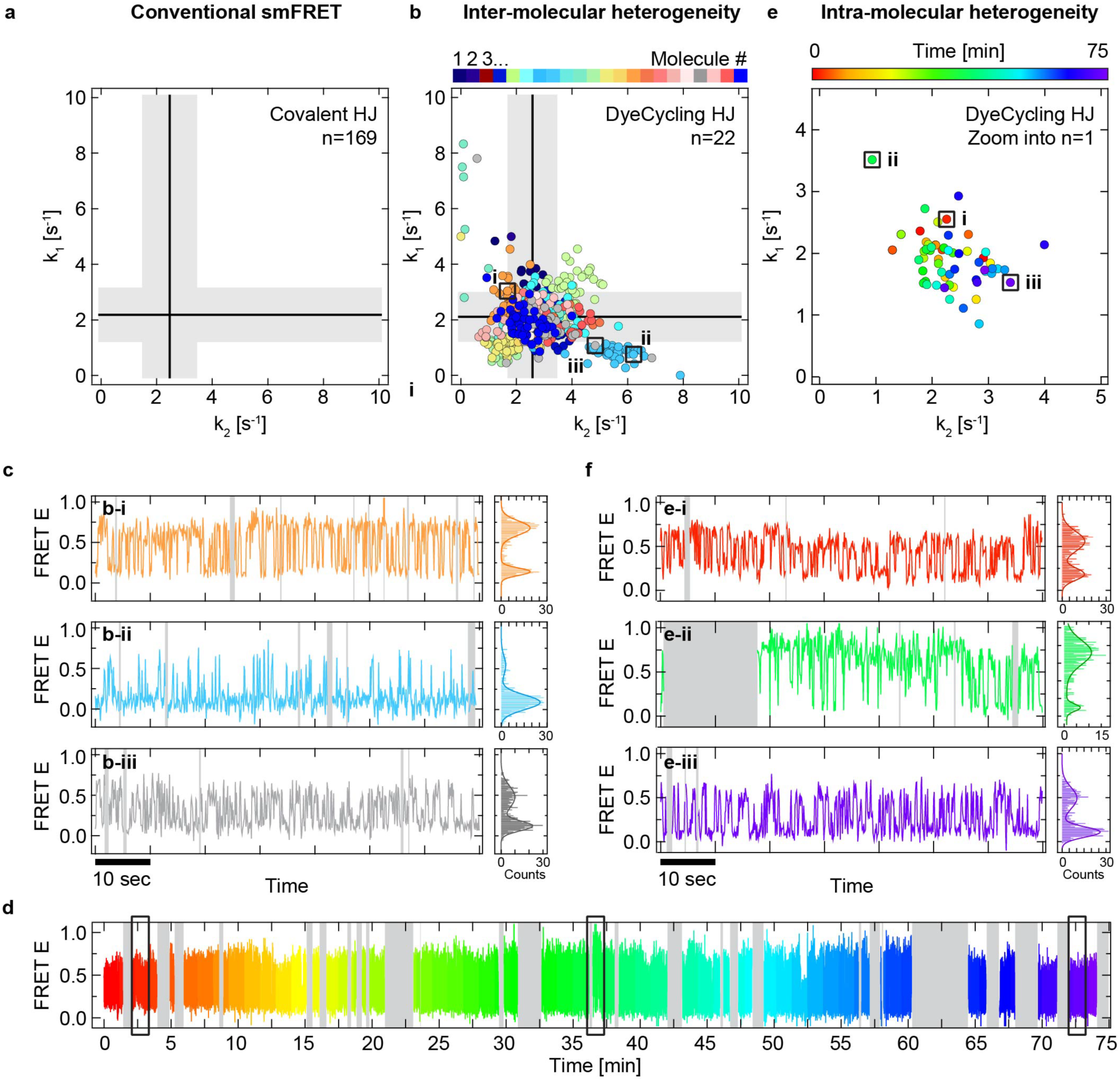
DyeCycling reveals inter- and intra-molecular heterogeneity. (**a**) Holliday junction (HJ) isomerization kinetics obtained by conventional smFRET: ensemble-averaged k_1_ and k_2_ (solid lines), standard deviation (grey shade). (**b**) Rate correlation plot showing the Holliday junction isomerization kinetics obtained by DyeCycling (realization 1). Each dot represents a k_1_-k_2_ pair extracted from a 70-second segment of a given DyeCycling trace, coloured per-molecule as indicated. Corresponding ensemble-averaged k_1_ and k_2_ (solid lines), standard deviation (grey shade), calculated after excluding the identified outlier (light blue cluster; see **Table S2** for details). The FRET efficiency traces of three labelled datapoints i, ii, iii) are displayed in (c). (**c**) Distinct kinetic behaviours observed in the FRET efficiency traces and histograms of three Holliday junction molecules (i, ii, iii), indicated in (b). (**d**) DyeCycling-enabled FRET efficiency trace of the Holliday junction molecule further explored in (e,f), coloured according to time with unbound gaps displayed in grey. (**e**) Rate correlation plot obtained from the single trace in (d) with rates extracted from consecutive 70-second segments, coloured according to time as indicated. The distribution of rates reveals the time-variant changes in the kinetics exhibited by a single Holliday junction molecule. The three framed rate pairs (i, ii, iii) specify the 70-second segments displayed in (f). (**f**) Distinct kinetic behaviours observed in the FRET efficiency traces and histograms of three different segments (i, ii, iii) of a single Holliday junction molecule, indicated in (e). (a-f) Sampling rate: 10 Hz (50 ms green, 50 ms red excitation). See methods for details. Covalent fluorophores used: Cy3B and ATTO647N. DyeCycling probes used: the Cy3B and ATTO643 probes (**Table 1**).

On the other hand, the per-trace kinetic analysis enabled by DyeCyling provides kinetic rates for each molecule, while ensemble-averaged values can be obtained for comparison; here k_1_ = 2.1 ± 0.9 s^-1^ and k_2_ = 2.6 ± 0.9 s^-1^ (**Fig. 4b**). The rate correlation plot (**Fig. 4b**) shows rate clusters colour-coded per molecule, where every datapoint represents a 70 s segment. Several rate clusters overlay with each other, suggesting they represent a single molecular species. The light blue cluster exhibits persistently distinct behaviour, with different kinetics and thermodynamics, indicative of static heterogeneity (see trace comparison in **Fig. 4c**). DyeCycling allows the analysis of this outlier separately to quantify its distinct rates (k_1_ = 5.5 ± 0.6 s^-1^ and k_2_ = 0.8 ± 0.2 s^-1^) and avoids pooling and averaging it together with the majority species. Noteworthy is also the grey dot labelled (iii) in **Fig. 4b**, which shows a temporary (70 s) distinct behaviour of a molecule exhibiting otherwise majority behaviour. This example of dynamic heterogeneity could be explained by a rare transition to a different kinetic regime arising from a high energy barrier crossing to different energy minima on the conformational energy landscape, or a distortion of the energy landscape possibly due to a rare molecular interaction. For Holliday junctions, particularly ionic interactions, viz. distinct Mg^2+^ coordination, have been found to induce such kinetic regime changes in Holliday junction conformational dynamics^64^. DyeCycling now provides the observation time to detect how these rare events occur spontaneously (see below). While more in-depth analysis is needed to fully uncover the molecular mechanism underlying Holliday junction behaviour, this example demonstrates that DyeCycling provides new means to investigate previously inaccessible kinetic effects. In particular, distinguishing conformational observations that originate from different molecular species from those representing temporary excursions on a single species’ energy landscape is highly valuable. By identifying such outliers, DyeCycling provides an important addition to the smFRET toolbox, preventing misinterpretation and enabling a more detailed understanding of the complex kinetics underlying biomolecular function.

Next, zooming into individual clusters allows one to assess kinetic heterogeneities within a single molecule. The DyeCycling trace in **Fig. 4d** and the corresponding rate correlation plot (**Fig. 4e**) are both color-coded for time (start to end of the trace). This molecule of the majority class (dark blue cluster in **Fig. 4b**) exhibits considerable dynamic heterogeneity and temporary thermodynamic inversion (k_1_/k_2_ ≈ k_2_/k_1_), which is evident in the three zoomed-in sub-segments plus histograms in **Fig. 4f**. While more advanced, pattern-specific segmentation would resolve the inherent heterogeneity more precisely, even the simple consecutive 70-s segments used here clearly reveal it. This “meandering” of the Holliday junction across the conformational energy landscape lacks a clear direction (red to blue in **Fig. 4f**), consistent with the absence of an external energy source in thermal equilibrium. Although distinct kinetic regimes of Holliday junctions have been observed previously by conventional smFRET^64^, the spontaneous transitions between these regimes and their order and timing, were previously undetectable. Thus, DyeCycling provides the observation time needed to capture these rare transitions and reveal the sequential order of distinct kinetic regimes displayed by a single molecule over periods ranging from several minutes to an hour (**Fig. 4**). While it is fascinating that molecules as simple as DNA Holliday junctions undergo such regime changes^64^, these kinetic changes were also found to play prominent roles in more complex systems, including RNA polymerase^24^, exonuclease^28^, and other biomolecular systems^29^. Collectively, these DNA-based studies demonstrate DyeCycling’s ability to (a) identify and investigate distinct molecular sub-species, rather than ensemble averaging them, and (b) provide detailed free-energy landscape sampling beyond the capabilities of bleaching-limited smFRET.

### DyeCycling of protein systems

Extending DyeCycling from DNA to protein systems is an important next step, because proteins often undergo complicated conformational dynamics with multiple states and rates that are difficult to fully capture and understand based on short, bleaching-limited smFRET traces. Longer DyeCycling-based smFRET traces could detect these dynamics more comprehensively and reveal the interrelation of different kinetic regimes observed in proteins. Conventionally, site-specific fluorescent labelling of proteins is commonly achieved by coupling maleimide-functionalized fluorophores to cysteine residues using the Michael addition. To this end, a single cysteine residue is introduced at the desired position by mutagenesis. For DyeCycling, we used the same approach to couple ssDNA oligomers to specific positions in the protein, which serve as docking strands for ssDNA-based fluorescent probes (**Fig. 5a**). As a test system for this proof-of-concept of protein DyeCycling, we chose the dimeric chaperone protein Hsp90 that is well characterized by smFRET, by us^33,49,66^ and others^32,67^. **Fig. 5a** shows the crystal structure of Hsp90’s closed conformation obtained under stabilization by a non-hydrolysable ATP analogue (adenylyl-imidodiphosphate, AMP-PNP)^68^. We coupled the docking strands to pre-validated positions^32^, indicated in **Fig. 5a**, via single cysteine point mutations in each of the Hsp90 monomers (see Methods). Nanophotonic protein DyeCycling with the pre-described self-healing fluorescent probes provided long, well-resolved smFRET observations. **Fig. 5b** shows a trace monitoring Hsp90’s conformational state for > 45 min with very good signal-to-noise ratio (zoom view in **Fig. 5b**). As expected, it detects predominantly a high-FRET state under saturating AMP-PNP conditions, consistent with the closed conformation shown by the crystal structure (**Fig. 5a**). The temporal coverage was 62% over the entire dataset which is comparable to DNA DyeCycling, whereas the reversible probe binding kinetics differ due to different buffer conditions.

**Figure 5.**
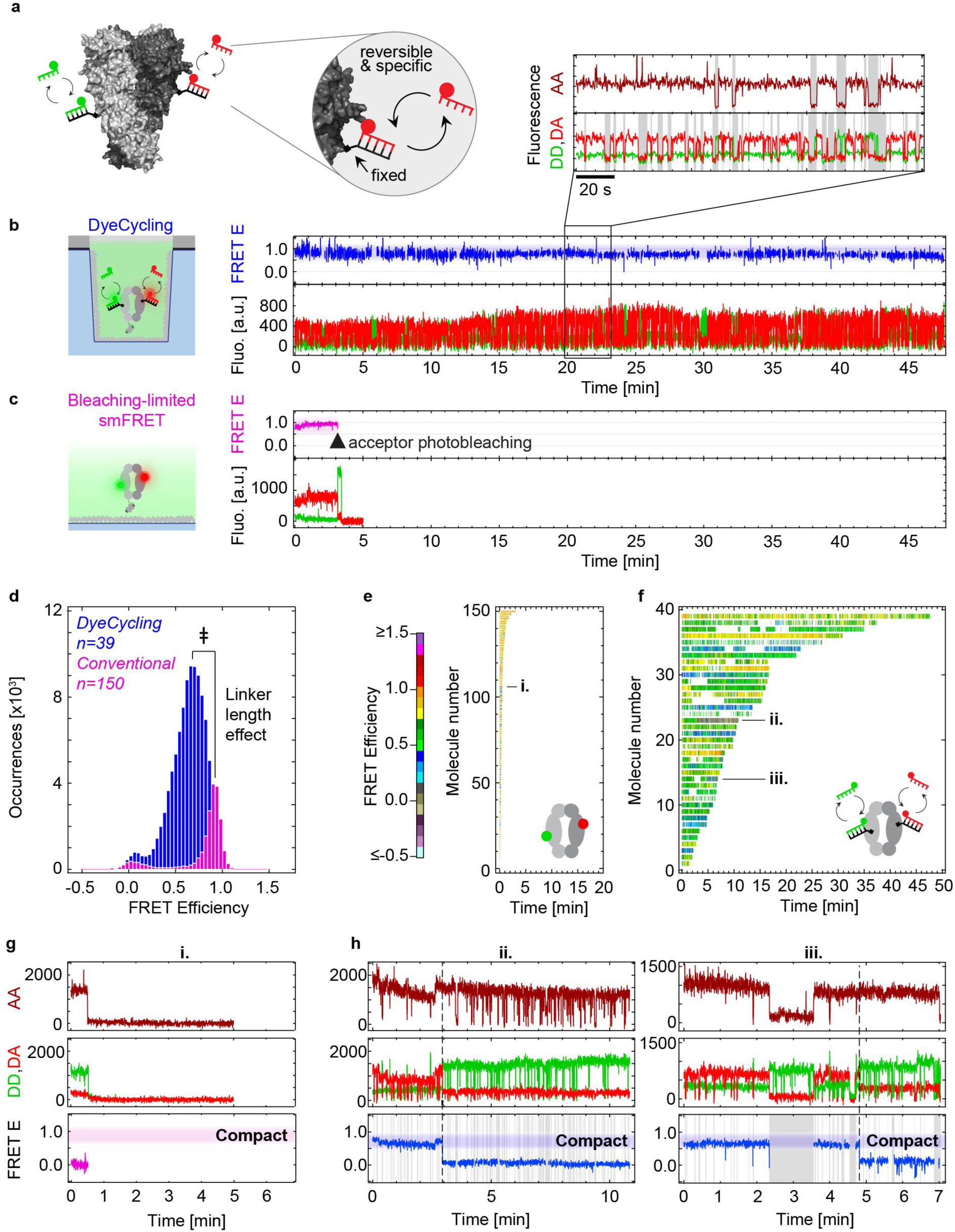
Protein DyeCycling proof of concept. (**a**) The DyeCycling protein construct: short ssDNA were site-specifically coupled to Hsp90, serving as docking strands for fluorescently labelled complementary ssDNA in solution. (**b**) Cartoon of the nanophotonic DyeCycling experiment (not to scale) and resulting DyeCycling trace of Hsp90 in the presence of 2 mM AMP-PNP. For over 45 min, the protein resides in the closed conformation. A zoom-view is included: DD, directly excited donor; DA, FRET-sensitised acceptor; AA, directly excited acceptor control; (partially) unbound gaps are shaded in grey. 5 Hz sampling (92 ms green, 92 ms red excitation) (**c**) The longest bleaching-limited smFRET trace of Hsp90 lasts for 3 minutes at identical 5 Hz sampling and scavenger conditions (see Methods). (**d**) Comparison of the FRET efficiency histograms obtained from 150 bleaching-limited smFRET traces vs. 39 DyeCycling traces, conditions as in (b,c). The double dagger ‡ indicates a peak shift caused by the longer fluorophore linker lengths in the DyeCycling experiment (for details see **Fig. S10**). (**e,f**) Whole-dataset plots of bleaching-limited smFRET (e) versus DyeCycling (f) with identical time axis scaling: all traces are colour-coded for FRET efficiency as indicated and sorted by trace length. e) Conventional smFRET provided 21’300 datapoints from n=150 molecules. (f) DyeCycling provided 94’638 FRET datapoints from 39 molecules. Unbound gaps appear white. (**g**) A rare low-FRET trace measured by bleaching-limited smFRET (conditions as in c). From the short observation, it remains unclear whether it represents a functionally relevant rare state or an artefact. (**h**) Protein DyeCycling resolves the ambiguity in (g): rare transitions between high-FRET and low-FRET states (i.e., compact to extended conformation) are indeed observed in the longer DyeCycling traces, indicating a bona fide effect rather than an artefact. (Partially) unbound gaps are shaded in grey. Covalent fluorophores used: Cy3B (N298C) and ATTO647N (A327C). Fluorescent probes used: LD555 and LD655 probes (**Table 1**) with docking strands at residues N298C and A327C, respectively. FRET windows shorter than 1 s were filtered out to exclude bias from non-specific probe adsorption.

In contrast to DyeCycling, bleaching-limited conventional smFRET yielded < 3 min observation time under the same conditions (5 Hz sampling, **Fig. 5c**). The FRET histograms (**Fig. 5d**) show prominent high-FRET populations in both cases as expected. Compared to the conventional smFRET high-FRET peak (E = 0.9 ± 0.065, standard deviation), the DyeCycling high-FRET peak is shifted and broader (E = 0.65 ± 0.16), which can be attributed to the longer linker between the fluorophore and the protein in this case (see **Fig. S10**). In addition, the histogram reflects again the significantly higher information content of DyeCycling traces, providing on average 2’426 FRET datapoints/molecule (13 min on average at 5 Hz sampling), compared to 142 datapoints/molecule for conventional, bleaching-limited smFRET traces (28 s on average at 5 Hz sampling). The information gain is also evident in **Fig. 5e,f**, providing a global comparison of conventional smFRET and protein DyeCycling, where each trace is colour-coded for FRET efficiency and sorted by trace length. The global yellow to green shift from Fig. 5e to 5f represents the mentioned fluorophore linker effect, cf. Fig. 5d. Overall, we demonstrated that nanophotonic DyeCycling is transferrable from DNA systems to protein systems, using the same established coupling chemistry and reversible probes. It provides long and well-resolved smFRET observations, offering more than an order of magnitude more data per single molecule.

Lastly, by zooming into the FRET traces in **Fig. 5g,h**, we demonstrate how the long DyeCyling traces can answer a question on the Hsp90 protein that was left unanswered by conventional smFRET. Note that Hsp90’s slow ATPase activity and its functional role remain debated in the field^69^, and nucleotide analogues, like AMP-PNP, are frequently used to study this matter^67,70^. Consistent with the Hsp90 crystal structure, conventional smFRET confirmed the compacting effect of AMP-PNP, leading to the closed conformation also in solution^67,71^. However, the origin of the small low-FRET population, indicative of an extended open conformation, was not fully understood. As an example, **Fig. 5g** shows a conventional smFRET trace showing a static low-FRET outlier among a majority of high-FRET observations. Is it representative of a different molecular species, possibly defective molecules? Or is it a rare state of fully functional Hsp90 protein? These questions remained unresolved based on the bleaching-limited observation fragments obtained by conventional smFRET. DyeCycling now provides compelling evidence that the low-FRET observations arise from defective Hsp90 proteins. First, DyeCycling reveals a smaller low-FRET population than bleaching-limited smFRET (**Fig. 5d**). Since DyeCycling enables more stringent trace selection – longer observations increase the likelihood of identifying artefacts – this result suggests that at least some of the low-FRET observations in the conventional dataset are artefactual. Second, the open state (low-FRET) population in the DyeCycling data arises from a minutes-long open state (**Fig. 5h**) out of which no open-to-closed transitions were detected in the entire dataset comprising 5.2 hours. However, the predominant closed population, consistently found under saturating AMP-PNP conditions, requires faster closing than opening rates. The apparent irreversibility from closed to open is therefore thermodynamically incompatible with Hsp90’s structural behaviour. Collectively, these findings suggest that the rare opening events represent denaturation of previously intact Hsp90, and the resulting low-FRET minority population arises most likely from damaged protein. More generally, this example shows how protein DyeCycling resolves uncertainties about trace-to-trace differences frequently observed by conventional smFRET, distinguishing whether they represent distinct molecular species (static heterogeneity, possibly impurities or damaged molecules) or fragmented observations of a single species due to photobleaching (dynamic heterogeneity, complex molecular behaviour).

Altogether, this proof-of-concept study (i) demonstrates the feasibility of protein DyeCycling, shown here using the complex multi-domain protein Hsp90; (ii) it provided long-term observations with high sensitivity and time coverage; and (iii) DyeCycling provided an explanation for a protein-specific question on Hsp90 that was left unsolved by conventional smFRET.

### Conclusion

Thirty years after the first demonstration of single-molecule FRET for studying biomolecules and their dynamics^72^, this work establishes nanophotonic DyeCycling to overcome smFRET’s longstanding photobleaching limitation. By harnessing the cumulative photon budgets of many reversibly bound fluorophores in sequence, DyeCyling experiments are not limited by single bleaching events and thus provide minutes-to-hour-long single-molecule observations – on average 60-fold more than conventional smFRET (**Fig. 3**). Nanoscale conformational changes are sensitively resolved and consistent with conventional smFRET. The background signal of fluorescent probes in solution is efficiently suppressed by zero-mode waveguides specifically developed for DyeCycling (**Fig. 2**). Compared to fluorogenic approaches^46^, this nanophotonic approach, applicable to DNA and protein systems, enabled better signal-to-noise ratio and ensures full flexibility in the choice of fluorophores.

By decoupling smFRET from photobleaching, DyeCycling adds essential capabilities to the smFRET framework. (i) Kinetic rate extraction at the single-molecule level enables new mechanistic and comparative insights (**Fig. 3**). (ii) Slow kinetic regime changes of single molecules are directly resolved, enabling the investigation of previously undetectable kinetic paths and their functional consequences (dynamic heterogeneity in **Fig. 4d-e**). (iii) Heterogenous observations arising from distinct molecular subpopulations, whether functionally relevant or artefactual, are individually identified enabling their separate investigation (**Fig. 4b,c** and **5g,h**). These conceptual advances on DNA and protein systems overcome several limitations of conventional smFRET, enabling deeper mechanistic insights and new types of smFRET experiments; for example, testing multiple physicochemical conditions sequentially on the same molecule, including biochemical interactors, temperature jumps, buffer conditions, etc. DyeCycling will allow researchers to systematically assess the significance of rare observations, such as long-lived enzymatic pausing^24^, frequently reported ‘static traces’^35–40^, and post-translationally modified subpopulations^29^ – questions that are often inaccessible to conventional, bleaching-limited smFRET. In the future, more direct and possibly externally triggered fluorophore coupling, enabled by unnatural amino acids and genetic code expansion^73^, could eliminate fluorophore labelling steps in the long-term. In conclusion, we anticipate that bypassing the photobleaching limit in smFRET will enable new dynamic insights and promote smFRET-based discoveries across the molecular life sciences.

## Materials and methods

All chemicals were purchased from Sigma-Aldrich, unless specified otherwise. All prepared solutions were filtered before use (0.22 μm nylon membrane by Merck, Germany).

### Palladium zero-mode waveguide fabrication

The specimen consisted of a 22 mm x 22 mm high-precision (1.5H) borosilicate coverslip (Paul Marienfeld, Germany) – pre-treated by burning at 500°C for 1 h, bathing in 2% Hellmanex III® in ultrapure water for 2 h, ultrapure water wash, nitrogen gas flow drying, and oxygen plasma treatment (5 min, 0.2 mbar, 30 W, 0.5 L/h oxygen gas flow) – coated with a Ti adhesion layer (5 nm) and Pd (95 nm) deposited via an electron beam evaporator (custom setup with components from Telemark (US), Inficon (CH) and Sharon Vacuum (US)). Focused ion beam (FIB) milling was performed on two instruments. For Fig. 2-4, a FEI Helios G4 CX system was used, equipped with a Ga ion source. The ion beam operated at an acceleration voltage of 30 kV, with beam currents adjusted between 24 pA to 40 pA depending on the desired ZMW size. The milling process was automated through the NanoBuilder software, setting the ZMW diameters to 50, 100, 150, and 200 nm. The pitch distance between individual ZMWs was set at 1 μm. Throughout the milling process, the chamber vacuum was maintained at 2 × 10^-6^ mbar. For Fig. 5, the FIB milling of over-milled ZMW arrays was performed using a Zeiss Crossbeam 540, equipped with a Ga ion source. An acceleration voltage of 30 kV and beam current of 50 pA was used. The process was automated using the SmartFIB software, setting a diameter of 65 nm, an ion dose of 80 to 90 mC/cm^2^ and a pitch of 1 μm to obtain holes with 100 nm diameter, milled through the metal layer (100 nm) and over-milled into the glass (150 to 175 nm). Cross-sections of a select few holes were cut using a 10 pA ion beam, to control the quality of the milling. The system vacuum during the milling process was maintained at 9.0 × 10^-8^ mbar.

### ZMW functionalization and passivation

Before each use, ZMWs were cleaned with acetone (on tissue, gentle scrubbing), ultrapure water, 10% household detergent solution (on tissue, gentle scrubbing), and ultrapure water again. Then the ZMW device was dried by gentle nitrogen gas flow, followed by oxygen plasma treatment (5 min, 70 millitorr, 100 W RF power using a Plasma Prep III, SPI supplies, USA), 20 min sonication in 10% detergent solution followed by three rounds of ultrapure water wash and 5 min sonication in ultrapure water, 20 min sonication in acetone, and drying by gentle nitrogen gas flow. ZMW were either stored under nitrogen gas at -20°C or immediately prepared for measurements. For functionalization, ZMW received oxygen plasma treatment (for wettability) and then were functionalized with Vectabond (3-aminosilane, Vector Labs, USA) at 1% in spectroscopy-grade acetone (Uvasol®). Measurement wells were created by sticking silicone gaskets with culture wells (Grace Biolabs, USA) onto the functionalized slide. The glass bottom of the ZMW was first passivated with a mixture of 11.2% (w/v) polyethylene glycol (mPEG-succinimidyl valerate MW 5,000, Laysan Bio Inc., USA) and 0.4% (w/v) biotinylated PEG (biotin-PEG-succinimidyl carbonate MW 5,000, Laysan Bio Inc., USA) in saturated K_2_SO_4_ buffer^74^ (0.1 mM sodium bicarbonate, 0.55 mM potassium sulphate, pH 9.5) for at least three hours at room temperature or overnight at 4°C while preventing dehydration in a humid chamber (pipet box). Gasket wells were washed with ultrapure water and dried with gentle nitrogen gas flow. In a second passivation round to optimally reduce potential fouling regions^75^, MS4-PEG (ThermoFisher, USA) at 250 µM in dimethyl sulfoxide (DMSO) was diluted tenfold in sodium bicarbonate buffer (0.1 M, pH 8.5) and incubated for ≥1h or overnight (in humid pipette tip box), then the gasket wells were washed with ultrapure water. Passivated ZMW were either immediately used, or stored under nitrogen gas at -20°C until use within three months.

### DNA Holliday junction design and annealing

The Holliday junction design was based on the HJ7 design described previously^63^ (**Table 1**). High-performance liquid chromatography (HPLC)-purified ssDNA strands (Biomers, Germany) were diluted to 1 μM in TN50 buffer (10 mM Tris-HCl, 50 mM NaCl, and pH 8) and annealed with a thermocycler (T-Gradient Thermoblock, Biometra, Germany) for 10 min at 90°C followed by cooling to 20°C (1°C·min^-1^).

#### Protein production

Yeast Hsp90 proteins with N298C (SUMO-yHsp90-N298C-Z-Avi) and A327C (yHsp90-A327C-Z-StrepII) cysteine mutations were produced as previously published^76^, with additionally 8 ml glycerol per litre expression media, and 1 mM TCEP in the anion exchange and size exclusion chromatography buffers. The N298C construct was treated with SUMO protease before anion exchange chromatography, as described^76^. The N298C construct was biotinylated during expression at a C-terminal AviTag; the A327C was not (no AviTag). For protein labelling, maleimide-functionalized ssDNA docking oligomers were purchased (Biomers, Ulm, Germany). Proteins were labelled as previously described for maleimide-functionalised dye attachment to cysteines^32^, with two-fold excess of maleimide functionalised fluorophores or ssDNA docking oligomers for DyeCycling, and sample clean-up by the gravity protocol of PD MiniTrap G-25 columns. N298C was labelled with Cy3B or donor docking strand. A327C was labelled with ATTO643 or acceptor docking strand. Labelling was confirmed by microvolume absorption spectrometry. N298C and A327C were mixed 1:9 for monomer exchange. Sample was stored at 4°C for a maximum of 6 days. Prior to measurements, the sample was centrifuged to pellet aggregates (19’000 rcf, room temperature, 3 minutes) before ≥45 min incubation with AMP-PNP.

#### Lumidyne probe preparation

Amine-functionalized ssDNA oligomers were purchased (Biomers, Ulm, Germany). NHS-LD555 and NHS-LD655 were purchased (Lumidyne Technologies, NY, USA). Lyophilized fluorophores were dissolved in anhydrous DMSO to 1 mM, and ssDNA oligomers were dissolved in potassium borate buffer (50 mM boric acid, 500 mM KCl, pH 8.1) to 1 mM. Fluorophores were conjugated to ssDNA by mixing ssDNA:potassium borate buffer:DMSO-dissolved fluorophores in ratios of 1:7.5:2.5, with a final DMSO content of 22.75%. The mixtures were incubated in the dark (250 rpm shaking, 20-25°C, 2 hours), and the reaction was quenched by adding 1/50 (v/v) of Tris acetate solution (1 M Tris, pH 7.5 using glacial acetic acid). Next, labelled probes were precipitated using sodium acetate (3 M sodium acetate, pH 6 using glacial acetic acid) in ratios 1:1:7:27 of reaction mix:sodium acetate:ultrapure H_2_O:ice-cold 100% ethanol for 15h at -20°C. Precipitated probes were pelleted (13000 rpm, 4°C, 1 hour). Supernatant was carefully removed to reduce free fluorophore contamination. Pellets were dissolved in 500 µl ammonium sulphate solution (1.7 M NH_4_SO_4_, 10 mM NH_4_OAc, pH 5.8). Probes were purified from unconjugated ssDNA and remaining free fluorophores using hydrophobic interaction chromatography on a HiTrap Phenyl Sepharose 1 ml column (Cytiva) equilibrated with previous ammonium sulphate solution and elution using a 0-100% gradient of elution buffer (10 mM NH_4_OAc, 4.5% methanol, pH 5.8) over 20 column volumes. Lumidyne probe fractions were collected and dialyzed against ultrapure water using the Pur-A-Lyzer (1 kDa) Midi dialysis kit (Sigma-Aldrich), for at least 5 hours per 800 µl sample to sufficiently remove ammonium sulphate before ethanol precipitation for probe reconcentration (1:8:27 ratios of 3M sodium acetate:dialysed sample:ice-cold 100% EtOH, for ∼15 h at -20°C, followed by centrifugation (13’000 rpm, 4°C, 1 hour)), pellet drying by nitrogen gas flow, and solvation in ultrapure water. Lumidyne probe concentration was estimated by absorbance measurements (DS11, Denovix, DE, USA), adjusted to 20 µM, and stored at 4°C until use. Initial labelling efficiencies were ∼80%; unlabelled ssDNA oligomers were removed during hydrophobic interaction chromatography. Recovery after each precipitation step was ∼50%. After purification, the fluorescent probes were near 100% pure (i.e., labelled and without contamination **Fig. S11**).

#### DyeCycling measurements

ZMW arrays were placed in the microscope objective focus, using markers for guidance. The passivated measurement well was incubated with 20 μL neutravidin (0.25 mg/mL, ThermoFisher, USA) for 15 min, washed with 600 μL start buffer (see below). For surface tethering of the unlabelled biomolecules, 20 μL sample (1-10 nM DNA or protein concentration) was added to the well for ∼1 min. Coverage of unlabelled biomolecules was estimated using 50 nM fluorescent probe solution diluted in start buffer, aiming for ∼80% of wells filled for optimal single-molecule occupation of 30-35% (according to Poisson statistics)^58^. The well was washed with ≥200 μL start buffer (for Hsp90 including 2 mM AMP-PNP), before imaging buffer was added. Acquisitions were 1 hour. Between acquisitions, imaging buffer was removed, the well was gently washed with start buffer, and fresh imaging buffer with probes was added. The ZMW slide was illuminated in TIR angle, which features stronger decrease of background signal as compared to wide-field excitation on ZMW^54^. The sampling rate was 10 Hz (50 ms green, 50 ms red excitation) for DNA DyeCycling (Fig. 3,4) and 5Hz sampling (92 ms green, 92 ms red excitation) for protein DyeCyling (Fig. 5).

For DNA Holliday junction experiments, these solutions were used: start buffer (50 mM Tris-HCl, 50 mM NaCl, 50 mM MgCl_2_, pH 8), fluorescent probe stock (20 µM probes coupled with Cy3B or ATTO643, or LD555 or LD655 (**Table 1**) in ultrapure water), freshly prepared imaging buffer (0.5-1 µM fluorescent probes, 1 mg/mL glucose oxidase, 0.04 mg/mL catalase (Roche Diagnostics, Switzerland)), and 1% (w/v) glucose for oxygen scavenging, 1 mM UV-matured Trolox^19^ in start buffer, saturated with nitrogen gas to mitigate radical oxygen-induced damage to DNA strands^65^).

For protein experiments, these solutions were used: start buffer (40 mM HEPES, 150 mM KCl, 10 mM MgCl_2_, pH 7.4), fluorescent probe stock (20 µM probes coupled with LD555 or LD655 fluorophore (**Table 1**), freshly prepared imaging buffer (1 µM fluorescent probes, 2 mM AMP-PNP, 3 U/ml pyranose oxidase, 90 U/ml catalase, 40 mM glucose, 2 mM cyclooctatetraene (Acros) and 0.5 mM UV-matured Trolox^20^ in start buffer saturated with nitrogen gas).

#### Coverslip control measurements

Regular coverslips for covalent control measurements were prepared as described previously^22^. Passivated coverslips were incubated for 5 minutes with NeutrAvidin, then washed with 600 µl start buffer. Covalently dye-labelled HJ or protein was diluted to picomolar concentrations for sparse surface-tethering to NeutrAvidin. Identical imaging buffers were used and identical laser powers were applied between control and DyeCycling, except for the replacement of fluorescent oligo solution with ultrapure water, and the exclusion of nitrogen saturation in the control HJ imaging buffer.

#### Microscopes

DNA and protein DyeCycling was measured on a custom-built TIRF setup with two Kinetix 22 cameras (s-CMOS, Teledyne Photometrics, AZ, USA) – one per spectral channel set by a dichroic mirror (ZT640rdc-UF3) and filters from Chroma, VT, USA: donor (550-615 nm, using ET585/65m), acceptor (655-755 nm, using ET706/95m). Additionally, a Nikon Eclipse Ti2-E microscope body with Perfect Focusing System was used with a Nikon CFI Apochromat TIRF 100x NA1.49 oil-immersion objective, a horizontal stage controller (MS-2000, Applied Scientific Imaging, USA), OBIS 532LS 150 mW and OBIS 637LX 160 mW lasers (Coherent, USA). Both lasers were used at 5.5 mW power (measured directly after the objective in epi mode), and TTL-triggered ALEX was controlled with a LabView script and a PCI card (PCIe-6535B connected by SHC68-C68-D4 to SMB-2163, National Instruments, USA). Early DNA DyeCycling data was acquired on a previously described TIRF setup^77^: EMCCD camera (iXon Ultra 897, Andor, UK), autofocus stabilization (CRISP, Applied Scientific Imaging, USA), a horizontal stage controller (MS-2000, Applied Scientific Imaging, USA), Arduino NANO V3-based (VNGSystems, NL) TTL triggering of lasers, using an ALEX time resolution of 100 milliseconds (50 ms exposure per frame) with a fibre-coupled 532 nm OBIS LS 150 mW laser (Coherent, USA) and 638 nm diode laser (Lasertack, Germany). Laser power was set at 10 mW (measured directly before the objective).

#### Data processing & analysis

See **Fig. S7** for data processing pipeline. Hour-long acquisitions, split into sub-movies for downstream processing, were pre-processed in Fiji (ImageJ) to correct for drift. Single-molecule traces were extracted from TIF movies in Igor Pro (Wavemetrics, Oregon, USA) by a custom script built upon software of the Hugel lab (Freiburg, Germany). In brief, intensity traces are extracted from each sub-movie and concatenated into the full-length single-molecule trace. Background correction was performed manually for each included DyeCycling molecule, using trace segments where neither fluorescent probe was bound. FRET E correction factors^78^ were determined using donor- or acceptor-only (segments of) traces. Cycler binding state assignment was performed using SMACKS^49^ by Hidden Markov modelling (HMM) with five states: (0) acceptor probe-only, (1) donor probe-only, (2) donor plus acceptor probe in low-FRET state, (3) donor plus acceptor probe in high-FRET state, and (4) no probes (see **Fig. S8**). For Holliday junction DyeCycling data, state assignment was manually evaluated to further filter out aberrant datapoints (e.g., an additional donor or acceptor signal due to non-specific adsorption close to a pixel containing the molecule of interest). For Protein DyeCycling data, a sixth HMM state was included to filter out such aberrant datapoints. Non-specific adsorption was low (quantified in ZMW using the Cy3B and ATTO643 probes on Hsp90 without DNA docking strands) and most (>96%) adsorption events lasted <1 sec. Hence, both Holliday junction and protein DyeCycling data was filtered to exclude all FRET-sensitive windows lasting <1 sec to prevent bias from non-specific adsorption events. For analysis, custom scripts were prepared (provided via Zenodo; see “Data, code, and material availability”).

## Supporting information

Supplementary Information Vermeer et al

## Acknowledgements

The authors thank Johannes Holbein, Dina Grohmann, Max Martin, Jerome Wenger, Srijayee Ghosh, Xiliang Yang for helpful discussions, Thorsten Hugel and Bianca Hermann for Hsp90 plasmids, and the Micro-Spectroscopy Research Facility at Wageningen University for early measurement time. This work was supported by the Swiss National Science Foundation via the NCCR Molecular Systems Engineering (MSE, grant no. 51NF–40–205608), by the University of Basel, and at an early stage by the Dutch Ministry of education (OCW). S.C. was supported by a Delft Technology Fellowship and an ERC StG grant (SIMPHONICS, Project No. 101041486).

## Data, code, and material availability

Single-molecule fluorescence intensity and efficiency trace files and custom scripts utilized for this study were deposited on Zenodo (DOI: 10.5281/zenodo.18040272). Acquired raw movies comprised in total over 1 TB of data, hence the intensity and efficiency trace files were deposited. Custom Igor scripts developed for this study were deposited on Zenodo. Pd ZMW devices are made available to academic users at cost price and subject to available capacity: contact S.S for more information.

## Author contributions

S.S. conceived the DyeCycling concept and designed experiments. B.V. designed and performed experiments, analysed data, and wrote the first draft. D.H.S. and S.C. designed and fabricated ZMWs. F.Z. and A.V. analysed, optimised, and fabricated ZMWs. B.V. and S.S. wrote the manuscript in consultation with all authors.

