## Supplementary Information Vermeer et al for "Nanophotonic DyeCycling delivers single-molecule FRET beyond photobleaching to identify heterogenous dynamics in DNA and protein systems"

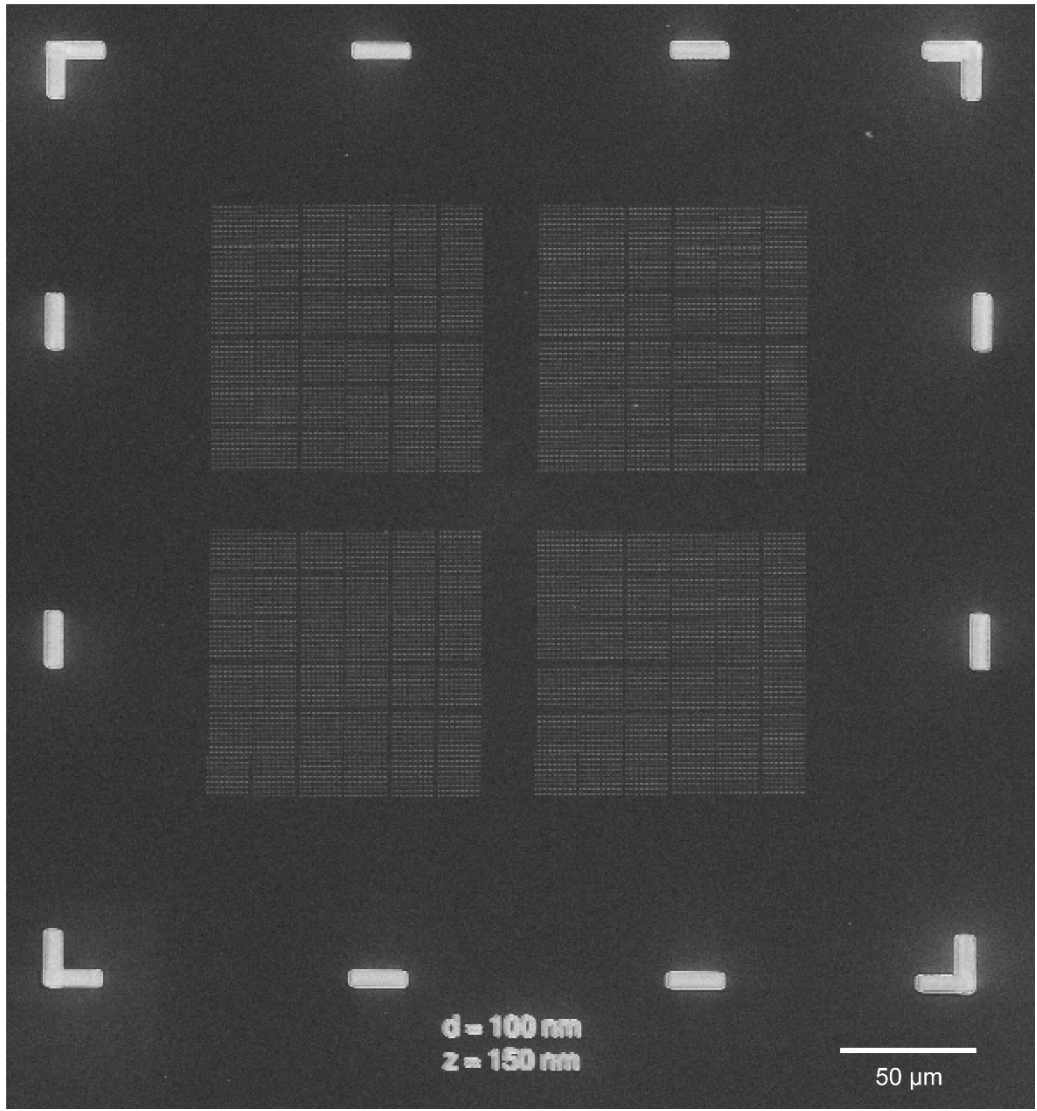

**Figure S1. Design of the zero-mode waveguide arrays.** Scanning electron micrograph showing the markers (outer edges) and ZMW arrays (inner area). The markers are visible by the naked eye, facilitating positioning the ZMW arrays under the microscope.

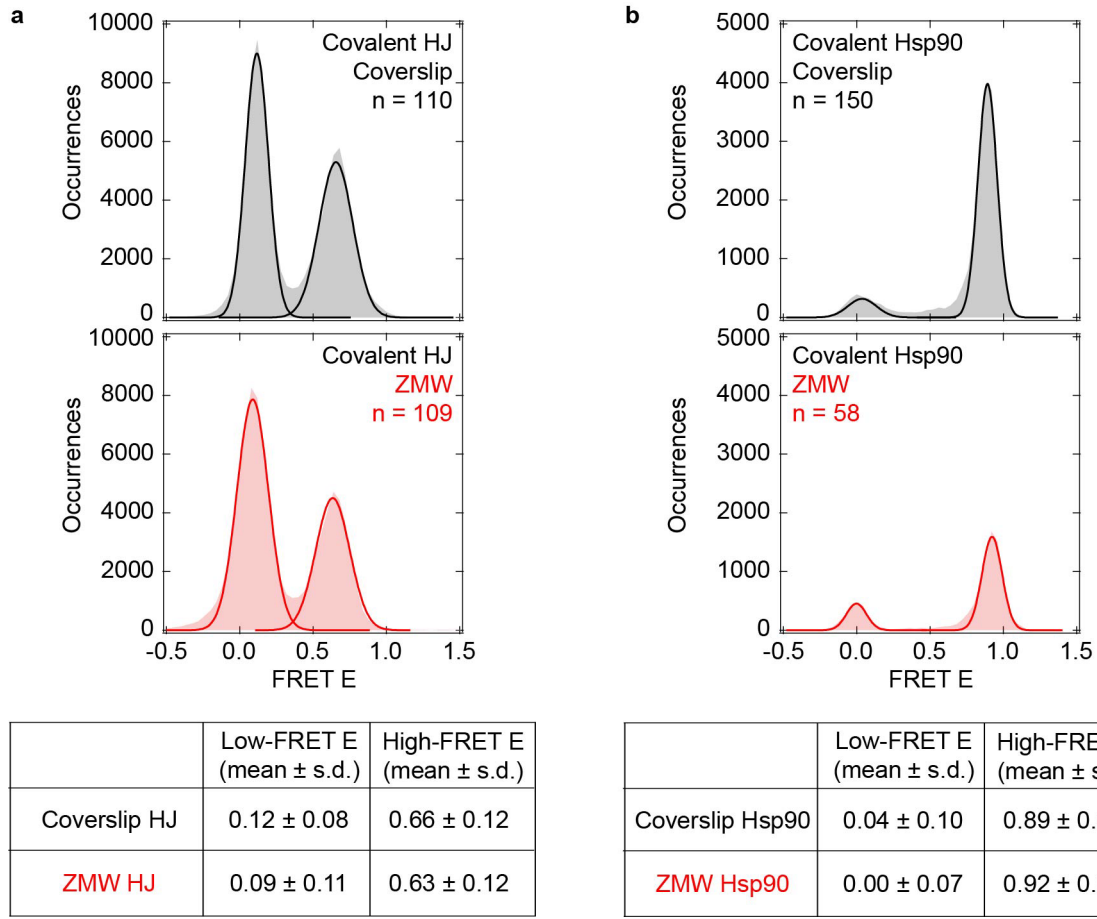

**Figure S2. FRET efficiencies measured on coverslips and in ZMW agree within experimental uncertainty.** (a) FRET Efficiency histograms obtained for covalently labelled Holliday junction (HJ) samples measured on a coverslip (black) and in ∅100 nm ZMWs (red). Gaussian fits are shown as solid line. The table shows the mean ± 1σ of the Gauss fit. Data were acquired on the sCMOS setup with 50 Hz sampling (7 ms green, 7 ms red excitation) in glucose oxidase scavenging imaging buffer (see Methods) with 200 mM MgCl<sub>2</sub>. Covalent fluorophores used: Cy3B and ATTO647N. (b) FRET efficiency histograms obtained for covalently labelled Hsp90 protein samples measured on a coverslip (black) and in ∅100nm ZMWs (red). Gaussian fits are shown as solid line. The table shows the population peak position ± standard deviation as derived by Gaussian fitting. Standard FRET corrections were applied (background, donor crosstalk, direct acceptor excitation, beta, and gamma)<sup>1</sup>. Data were acquired on the sCMOS setup with 5 Hz sampling (92 ms green, 92 ms red excitation) in pyranose oxidase scavenging imaging buffer with 2 mM AMP-PNP. Covalent fluorophores used: Cy3B (N298C) and ATTO647N (A327C).

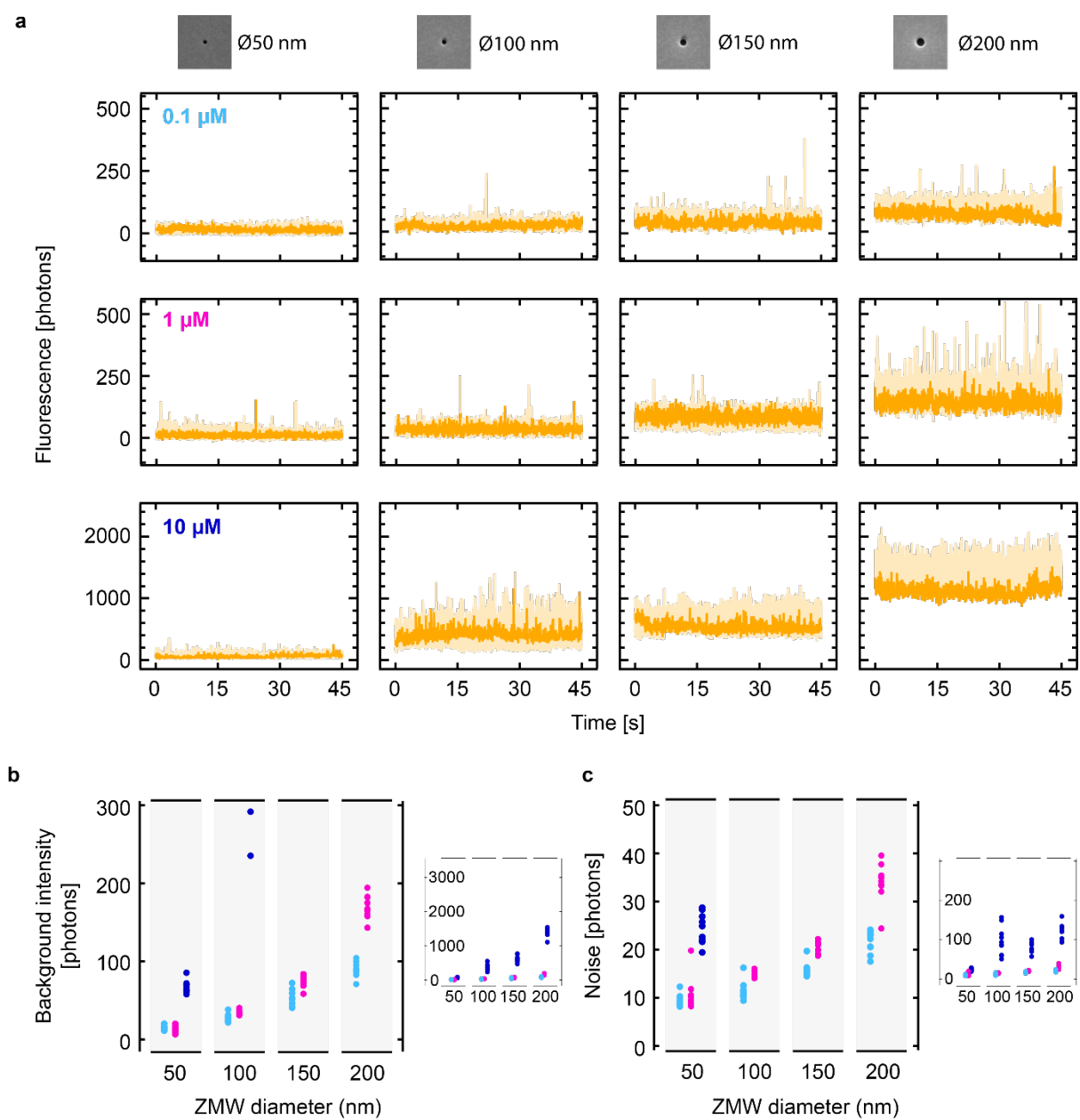

**Figure S3. ZMW diameter comparison: donor background and noise at increasing donor probe concentrations.** (a) Representative traces of background fluorescence induced by three different fluorescent donor probe concentrations (Cy3B probe, see main text **Table 1**), measured in individual, passivated waveguides of various diameters (10 traces are overlaid, with one trace coloured darker). No molecules were immobilized in the waveguides. (b,c) Quantification of data in (a). (b) Mean background intensity per trace ($n=10$ per condition). Right panel: zoomed-out view showing the values measured at $10\ \mu\text{M}$ donor probe. (c) Mean background noise per trace (standard deviations, $n=10$ per condition). Data were acquired on the EMCCD setup with 10 Hz sampling (50 ms green, 50 ms red excitation) in glucose oxidase scavenging imaging buffer (see Methods).

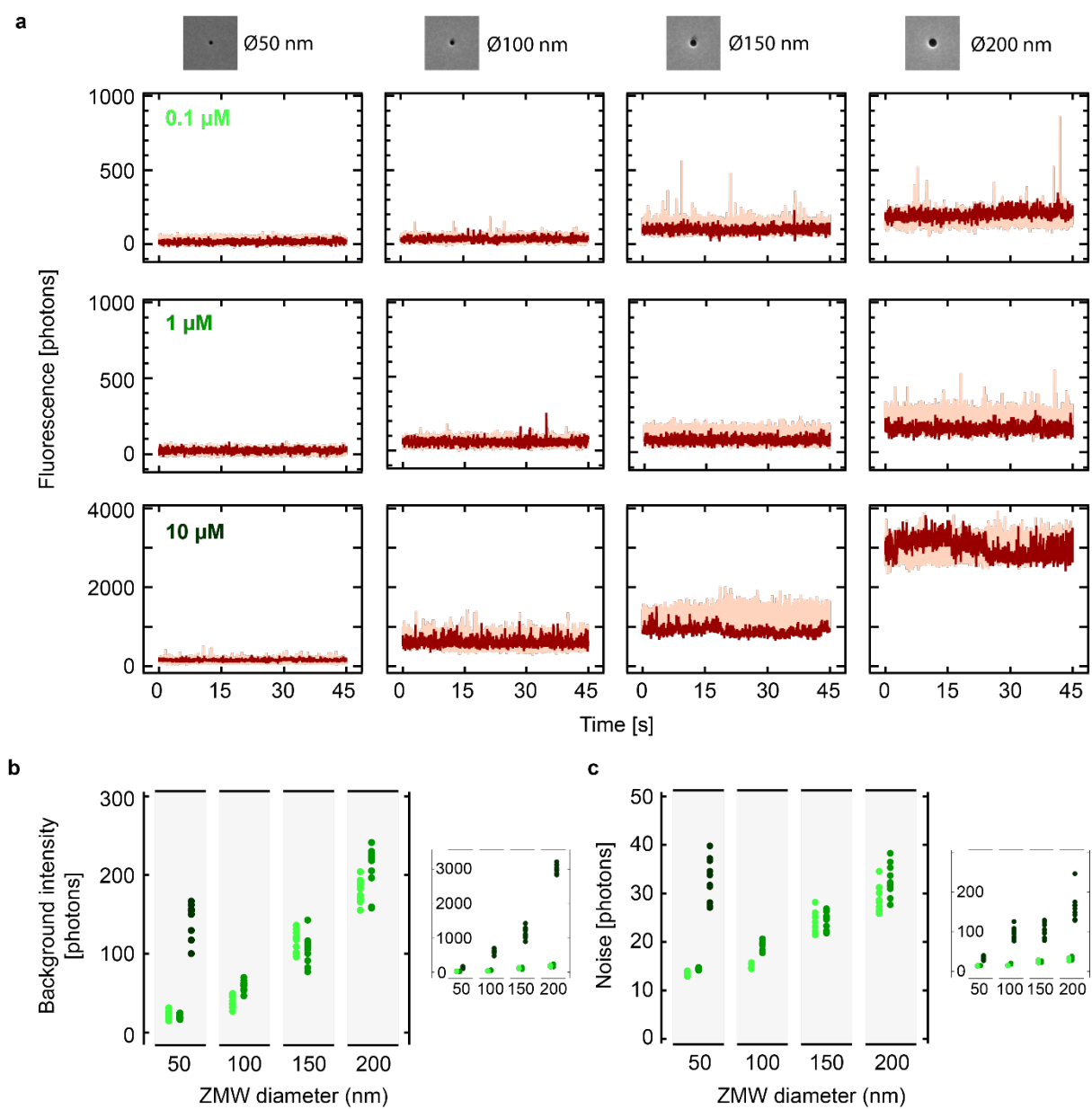

**Figure S4. ZMW diameter comparison: acceptor background and noise at increasing acceptor probe concentrations.** (a) Representative traces of background fluorescence induced by three different fluorescent acceptor probe concentrations (ATTO643 probe, see main text **Table 1**), measured in individual, passivated waveguides of various diameters (10 traces are overlaid, with one trace coloured darker). No molecules were immobilized in the waveguides. (b,c) Quantification of data in (a). (b) Mean background intensity per trace ($n=10$ per condition). Right panel: zoomed-out view showing the values measured at $10\ \mu\text{M}$ donor probe. (c) Mean background noise per trace (standard deviations, $n=10$ per condition). Data were acquired on the EMCCD setup with 10 Hz sampling (50 ms green, 50 ms red excitation) in glucose oxidase scavenging imaging buffer (see Methods).

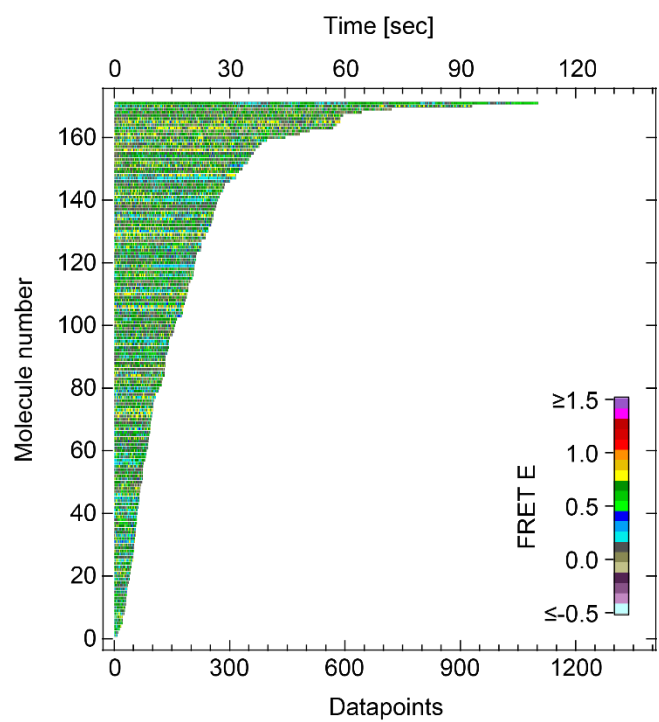

**Figure S5. Whole-dataset plot of Holliday junction data measured by bleaching-limited smFRET.** Traces are sorted by length and colour-coded by FRET Efficiency (FRET E), as indicated by the legend. Data were acquired on the EMCCD setup with 10 Hz sampling (50 ms green, 50 ms red excitation) in glucose oxidase scavenging imaging buffer (see Methods). Covalently attached fluorophores: Cy3B and ATTO647N.

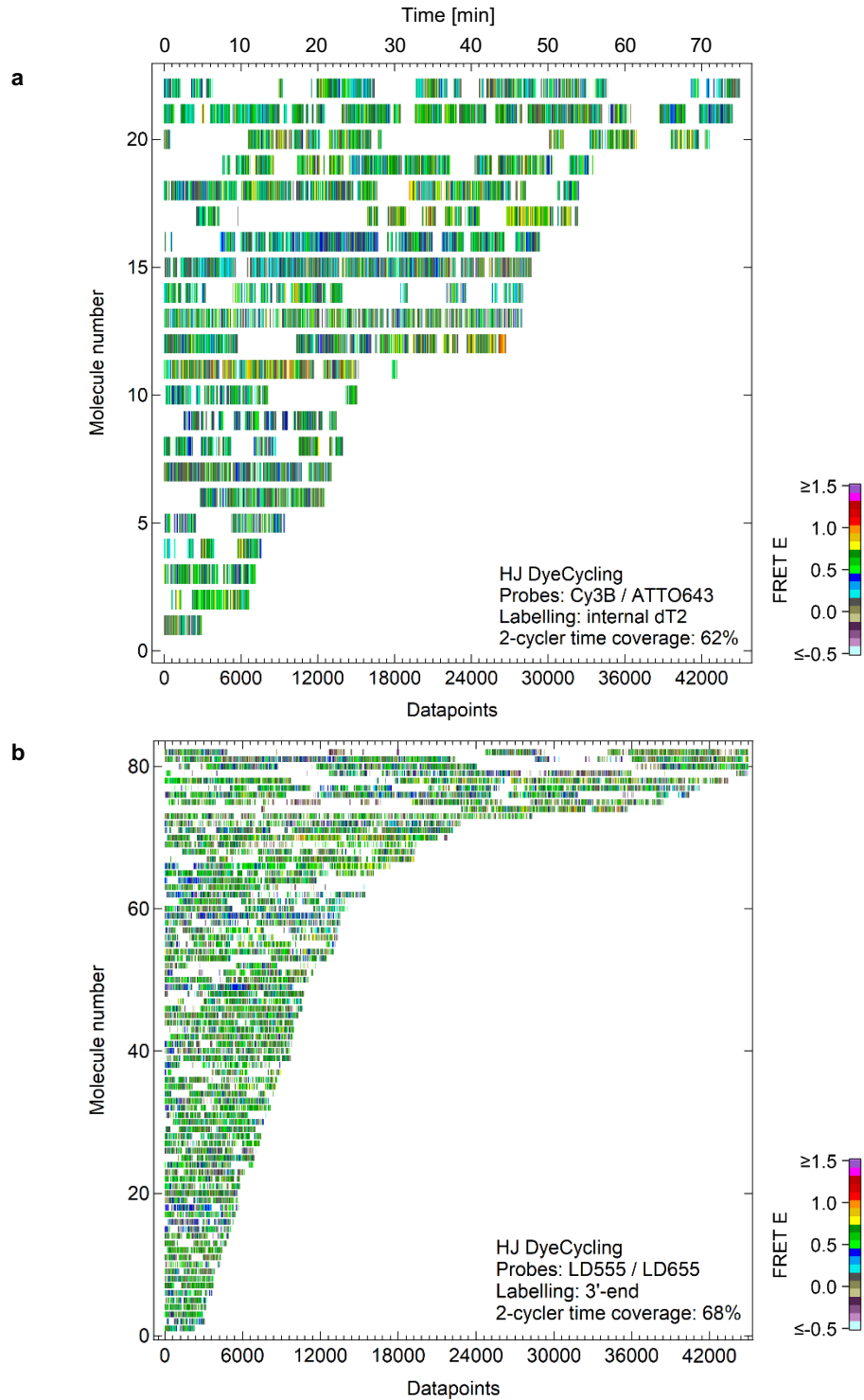

**Figure S6. Whole-dataset plot of DNA DyeCycling data measured for two different FRET pairs.** Traces are sorted by length and colour-coded by FRET Efficiency, as indicated by the legend. **(a)** Holliday junction DyeCycling data ($n=22$). Donor probe: Cy3B probe. Acceptor probe: ATTO643 probe (see main text **Table 1**). Probe concentrations were 500-750 nM. Data were acquired on the EMCCD setup with 10 Hz sampling (50 ms green, 50 ms red excitation) in glucose oxidase scavenging imaging buffer with 50 mM $MgCl_2$ (see Methods). **(b)** Holliday junction DyeCycling data ($n=82$). Same time axis as in (a). Donor probe: 3'-end-labelled with Lumidyne555. Acceptor probe: 3'-end-labelled with Lumidyne655 (see main text **Table 1**). Probe concentrations were 1 μM. Data were acquired on the sCMOS setup with 10 Hz sampling (50 ms green, 50 ms red excitation) in pyranose oxidase scavenging imaging buffer with 200 mM $MgCl_2$ (see Methods).

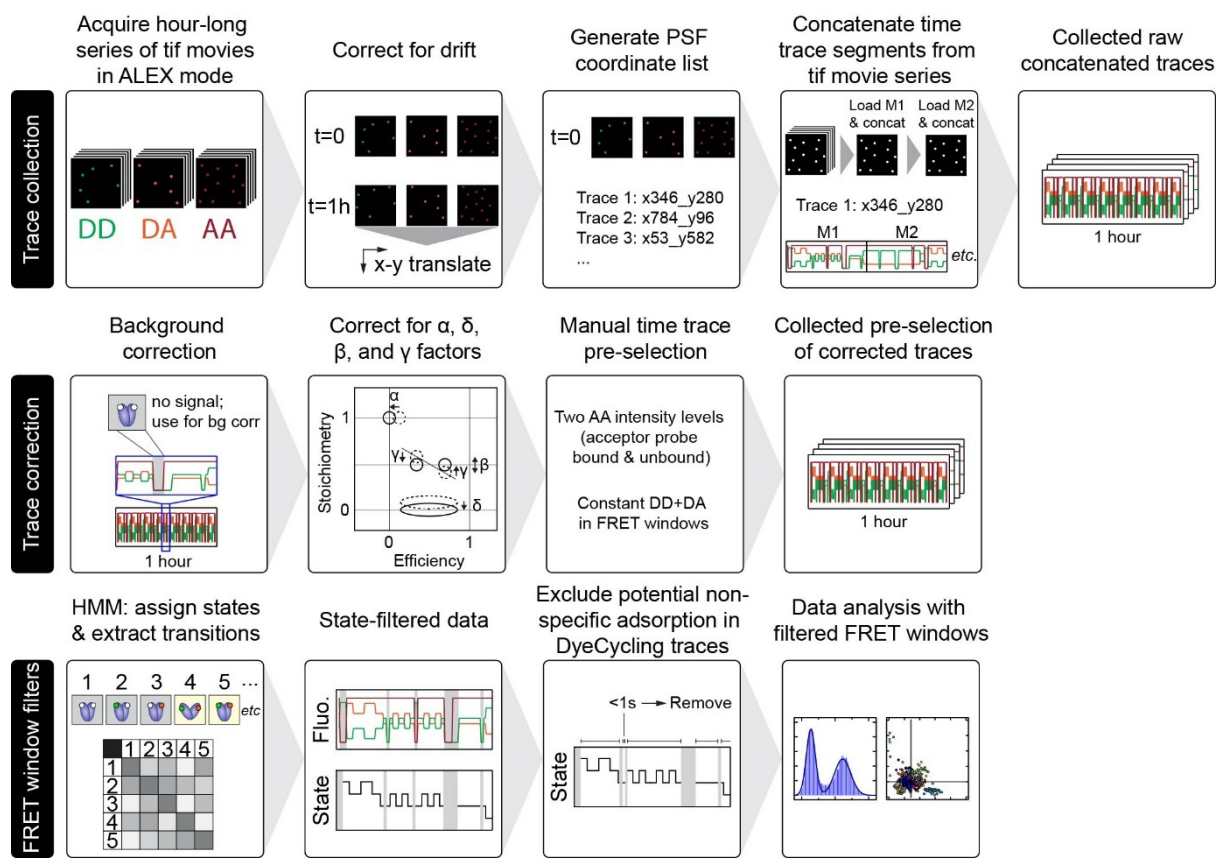

**Figure S7. Pipeline for data acquisition and processing.** **Trace collection:** Movie acquisition was performed with the Beacon software (Photometrics). Translation-based drift correction was performed with ImageJ's Translate functionality using drift tables compiled from the hour-long acquisitions. Point-spread function (PSF)-based single-molecule time trace extraction was performed through software developed for Igor Pro (Wavemetrics) by the Hugel lab. Large hour-long movie files (ca. 30-100 Gb) were loaded with a custom script for coordinate-based time trace concatenation through iterative sub-movie loading. This yielded a collection of raw concatenated fluorescence time traces. **Trace correction:** ZMW traces were corrected for background using trace segments where no donor and no acceptor signal was present. Leakage ($\alpha$), direct acceptor excitation ($\delta$), beta ($\beta$), and gamma ($\gamma$) factors were corrected as per smFRET field standards<sup>1</sup>. A manual pre-selection was made based on these criteria (i) constant directly-excited acceptor emission (AA) where present, (ii) a constant sum of donor-excited donor emission (DD) and donor-excited acceptor emission (DA) in the FRET windows. This yielded a collection of pre-selected corrected fluorescence and FRET efficiency traces. **FRET window filters:** FRET windows and gaps with (partially) unbound probes were assigned by Hidden Markov modelling using the SMACKS software<sup>2</sup> developed for Igor Pro. See Fig. S8. Only the FRET windows entered further analysis. In addition, all FRET windows shorter than 1 second were excluded from data analysis, to prevent potential bias from non-specific adsorption of fluorescent probes.

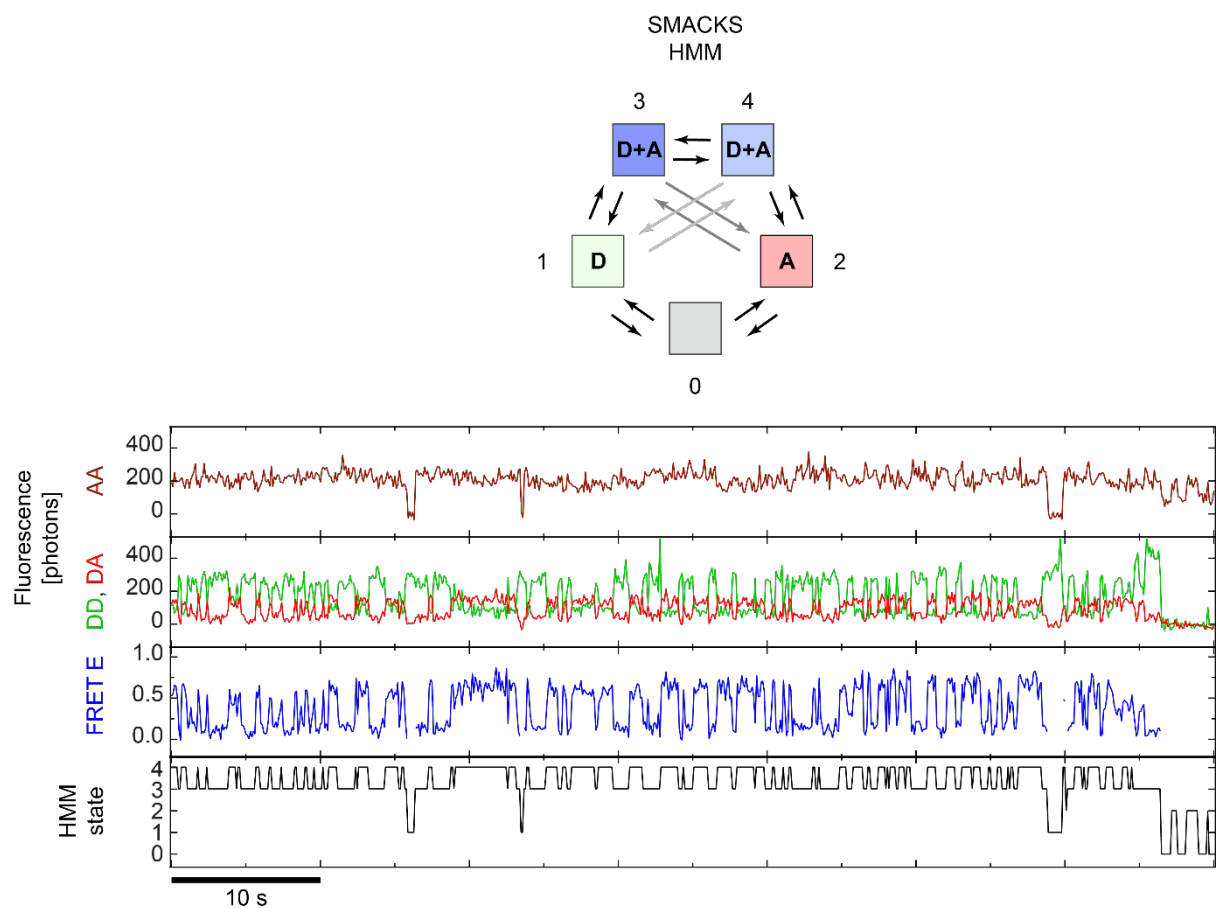

**Figure S8. Reliable DyeCycling gap identification by Hidden Markov modelling.** Kinetic scheme of the states occurring in a Holliday junction experiment: 0) acceptor only, 1) donor only, 4) completely unbound, 2,3) bona fide states, i.e. different conformations (number depends on the system under study). Using alternating laser excitation (ALEX), all these states can be distinguished. Hidden Markov modelling (HMM) was employed to identify all states using the SMACKS software<sup>2</sup>. For the Protein DyeCycling analysis, a 6<sup>th</sup> state was added to filter out aberrant datapoints, e.g., an additional donor or acceptor signal due to non-specific adsorption close to pixel containing the molecule of interest. Data were acquired on the EMCCD setup with 10 Hz sampling (50 ms green, 50 ms red excitation) in glucose oxidase scavenging imaging buffer (see Methods). Fluorescent probes used: Cy3B and ATTO643 probes (see main text **Table 1**).

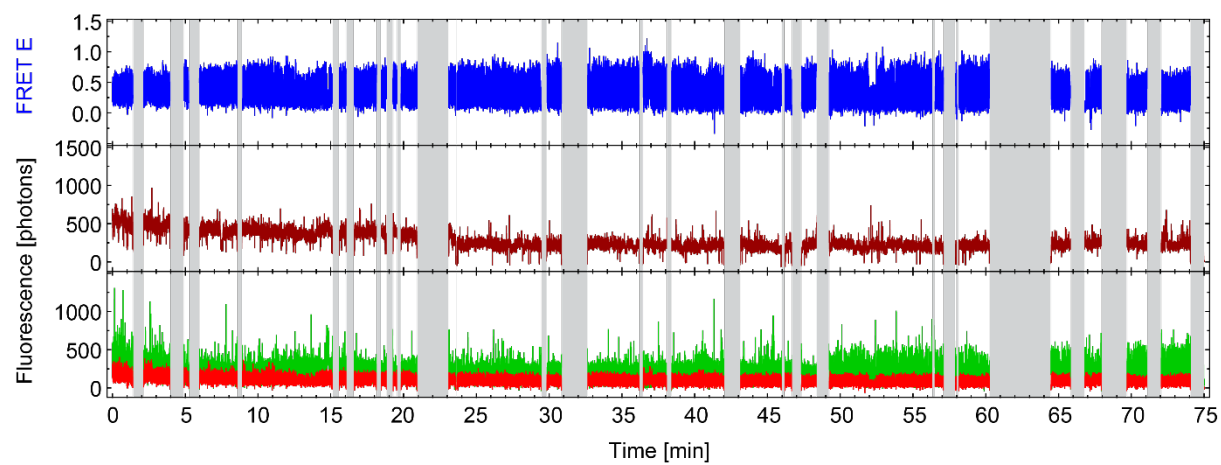

**Figure S9. Raw data accompanying main text Figure 3.** Holliday junction DyeCycling data with Cy3B donor probe and ATTO643 acceptor probe. Blue: FRET Efficiency; Dark red: directly excited acceptor fluorescence; Green: donor fluorescence; Red: FRET-sensitized acceptor fluorescence; Grey shade: blind gaps. Data were acquired on the EMCCD setup with 10 Hz sampling (50 ms green, 50 ms red excitation) in glucose oxidase scavenging imaging buffer (see Methods).

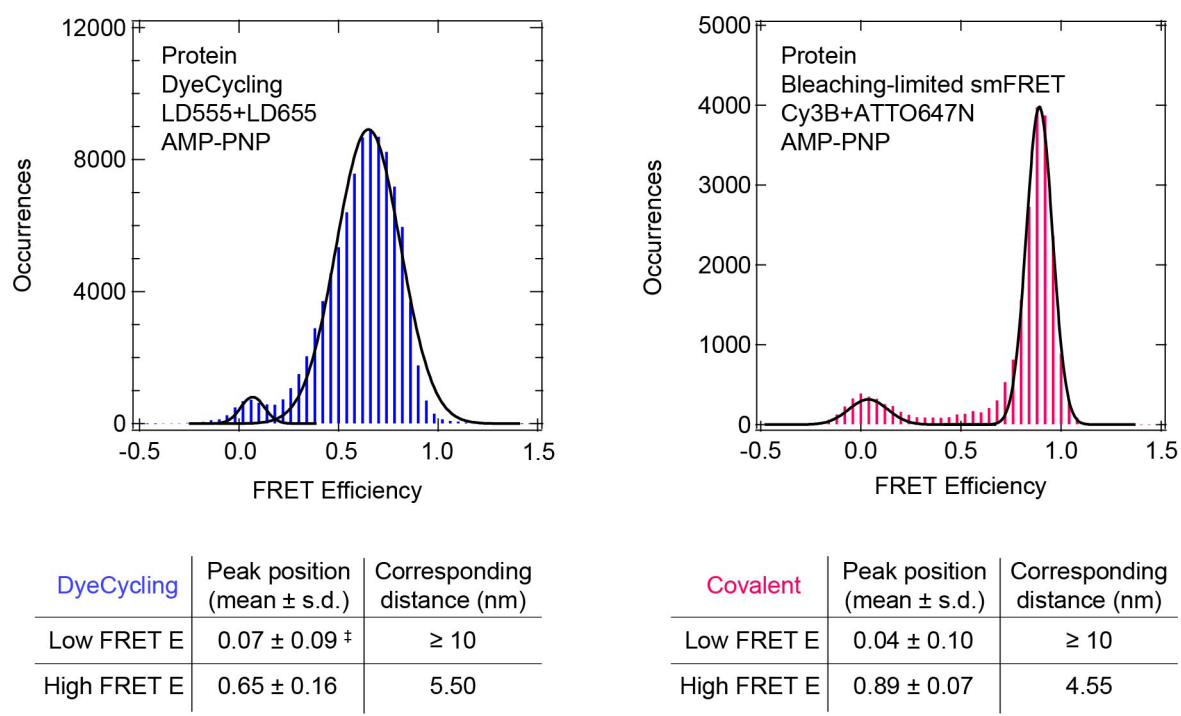

**Figure S10. Effect of fluorophore linker length on FRET efficiencies: protein DyeCycling vs. conventional smFRET with covalent labels.** FRET Efficiency histograms for DyeCycling (left, $n=39$) and conventional smFRET traces (right, $n=150$) with standard corrections applied (background, donor crosstalk, direct acceptor excitation, beta, and gamma)<sup>1</sup>. The table lists the mean FRET efficiencies obtained by Gaussian fits ($\mu \pm 1\sigma$) and estimated inter-dye distances. The longer linker lengths in DyeCycling experiments lead to a broader distribution and 1 nm longer inter-dye distances. The Förster radius for the DyeCycling pair (LD555, LD655) was $R_0 = 6.07$<sup>3</sup> and for the covalently coupled pair (Cy3B, Atto647N) $R_0 = 6.463$<sup>4</sup>. ‡ The DyeCycling low-FRET peak was fit with fixed width; all other coefficients were free. Data were acquired on the sCMOS setup with 5 Hz sampling (92 ms green, 92 ms red excitation) in pyranose oxidase scavenging imaging buffer with 2 mM AMP-PNP.

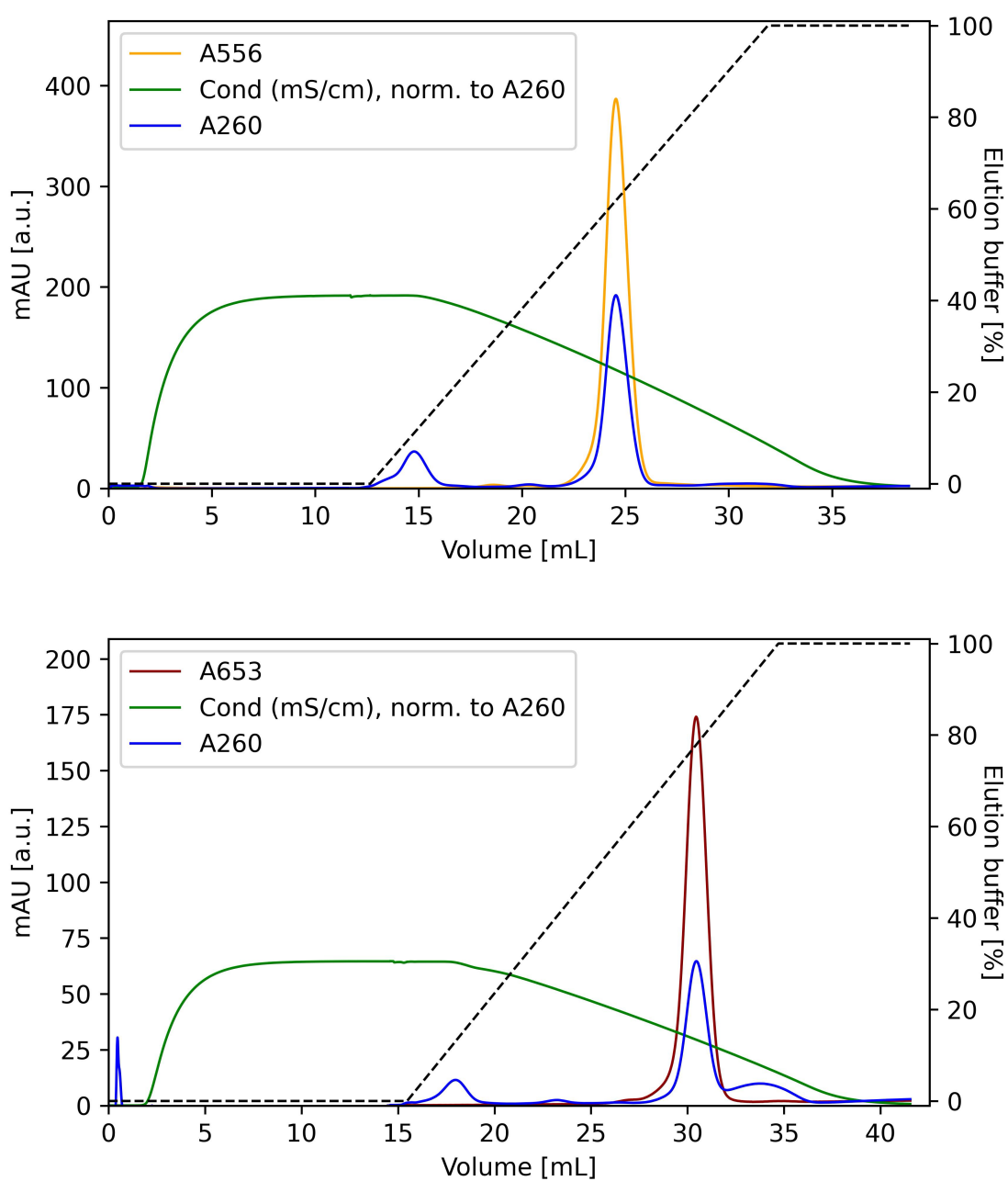

**Figure S11. Hydrophobic interaction purification of Lumidyne probes.** LD555- (Top) and LD655-labeled (Bottom) probes were separated from free DNA oligos and free fluorophores after the conjugation reaction. The elution buffer gradient (dotted line) caused elution of free oligos (first A260 peak), and fluorophore-labelled oligos appeared at ~65% (LD555 probe) and ~80% (LD655 probe) elution buffer. Probes were purified on a 1 ml Phenyl Sepharose column with a gradient from ammonium sulphate buffer (1.7 M $\text{NH}_4\text{SO}_4$, 10 mM $\text{NH}_4\text{OAc}$, pH 5.8) to elution buffer (10 mM $\text{NH}_4\text{OAc}$, 4.5% methanol, pH 5.8).

**Table S1. SmFRET good trace yield in coverslips versus zero-mode waveguides.** Percentages reflect the number of good traces of covalently labelled biomolecules among the total number of fluorescent spots that feature both donor and acceptor signal.

| **Sample** | **Coverslip** | **Zero-mode waveguide** |
| --- | --- | --- |
| Holliday Junction | 18% | 8.3% |
| Heat shock protein 90 | 4.5% | 1.5% |

**Table S2. Ensemble-averaged Holliday junction interconversion rates corresponding to data presented in Figure 4a,b.** Rates extracted by Hidden Markov modelling (SMACKS) were weighted by datapoints.

| **Condition** | ***k*1 mean ± s.d. [Hz]** | ***k*2 mean ± s.d. [Hz]** |
| --- | --- | --- |
| Covalent-labelled HJ (*n* = 169) | 2.2 ± 1.0 | 2.5 ± 1.0 |
| DyeCycling HJ, full dataset (*n* = 22) | 2.0 ± 0.9 | 2.9 ± 1.2 |
| DC HJ, excl. light-blue outlier (*n* = 21) | 2.1 ± 0.9 | 2.6 ± 0.9 |
| DC HJ, light blue outlier only (*n* = 1) | 0.8 ± 0.2 | 5.5 ± 0.6 |
